# Distinct signaling center and progenitor identity dynamics initiate human forebrain patterning

**DOI:** 10.64898/2026.03.25.714146

**Authors:** Afnan Azizi, Dana Fakhreddine, Fursham Hamid, Zeno Messi, Stephanie Strohbuecker, Francois Guillemot, Corinne Houart

## Abstract

At which developmental stage does the human brain start to differ from other mammals? We uncover human features of telencephalic patterning as early as the fourth post-conception week, comparing the early mouse and human forebrains using single-cell transcriptomics and 3D spatial multi-transcript imaging. In comparison to mice, the human telencephalon is delayed in ventral SHH signaling, accompanied by a human signature of anterior FGF signaling. These observations correlate with a less-resolved patterning and reduced progenitor diversity along the dorsoventral and anteroposterior axes, as well as a human early neurogenic signature. Our complementary approaches reveal a human divergence in allocation of telencephalic progenitor identities, propelled by temporal and qualitative differences in ventro-anterior signals following neural tube closure in the developing forebrain.

## Introduction

The human neocortex has evolved to be proportionally bigger and more complex than those of most other mammals. Recent studies have shown that a main mechanism in generating this complexity is spatio-temporal control of neurogenesis (1). Progenitors generate neurons after acquiring their anteroposterior and dorsoventral identities, a process called patterning (2, 3). While the overall mechanisms are conserved in mammals, relative sizes of the various progenitor territories across these organisms vary, suggesting evolutionary tweaks in the patterning process (4). It is, therefore, reasonable to assume that the differences in neurogenic behaviors between species are determined by differences in progenitor identities. Alas, the dearth of data from human embryonic samples at these stages has been a major obstacle in extending our knowledge to human.

Expansion of the neocortex in human has been primarily attributed to the diversification and proliferation of basal intermediate progenitors in the cortical subventricular zone that occur during the fetal stages of development (5–8). Comparatively, little attention has been paid to the earlier establishment of regional identities that give rise to the diversity of fates of the later cortex.

The neocortex arises from the embryonic telencephalon, where projection neurons are primarily generated in the dorsal part (pallium), and interneurons originate in the ventral part (subpallium) (9). The telencephalon is regionalized into pallium and subpallium through the action of morphogens and signaling pathways—i.e. FGFs anteriorly, SHH ventrally, and WNTs dorsally (Figure1B)—that are conserved in all vertebrates (10–12). However, they can differ in the timing of their activity. In fact, evidence of evolutionary modulation of various tissues and organs, including the telencephalon, through heterochronic and heterometric changes in morphogenic factors is accumulating (13–16). For instance, by comparing closely related cichlid fishes of lake Malawi, we and our colleagues demonstrated that the species expressing Shh earlier and more anteriorly had a larger subpallium than the one with a later, more posterior Shh expression. Indeed, manipulating Shh expression in the cichlid fish and zebrafish confirmed the hypothesis that higher levels of Shh leads to a larger telencephalon, with a bigger subpallium (17, 18).

**Fig. 1.**
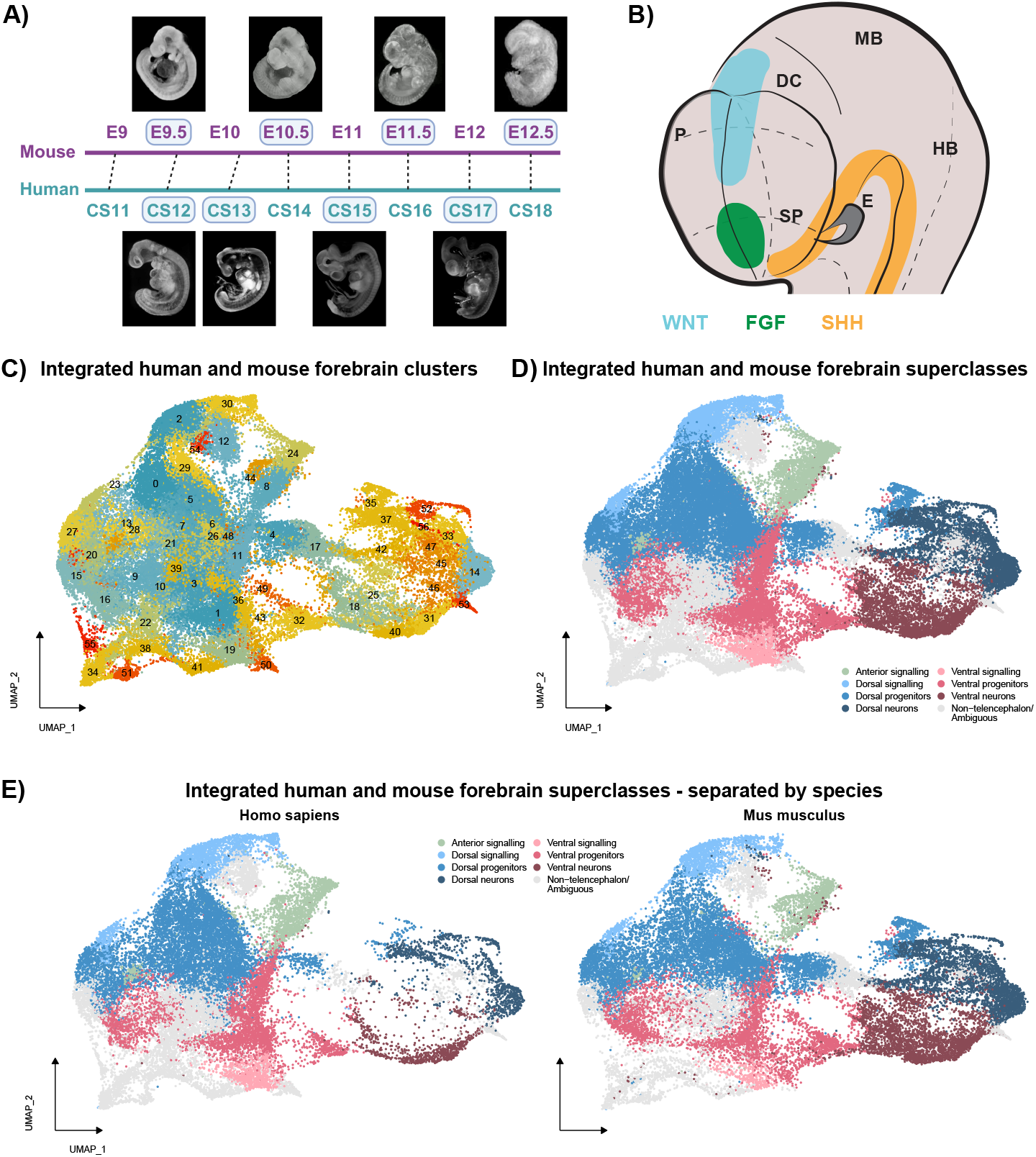
Single cell RNA sequencing of human and mouse embryonic telencephalon. **(A)** Timeline equivalency of mouse and human embryonic development as reported in the literature. The four stages used for each species are shown with images attached. Mouse embryonic images were obtained from the EMAP eMouse Atlas Project (http://www.emouseatlas.org) (21), and human embryonic images were obtained from the HDBR Atlas (https://hdbratlas.org/) (22, 23). **(B)** A schematic of the mammalian telencephalon at the phylotypic stage of development highlighting the location of the three main signaling centers patterning it. P: pallium; SP: subpallium; DC: diencephalon; E: eye; MB: midbrain; HB: hindbrain. **(C)** UMAP plot of the integrated human and mouse forebrain cells from the eight stages indicated in **(A). (D)** UMAP plot of the integrated human and mouse forebrain cells annotated with major cell type classes (superclasses). Anterior signaling (green), dorsal signaling (light blue), dorsal progenitors (cerulean), dorsal neurons (navy blue), ventral signaling (salmon pink), ventral progenitors (deep pink), ventral neurons (maroon), and other cells posterior to the telencephalon or unidentifiable (grey) are shown. **(E)** UMAP plots of integrated human and mouse forebrain superclasses separated by species.

Yet, many studies comparing human and mouse (or other species) development rely on stage equivalencies based on gross morphological characteristics, such as rump to crown length, limb growth, or cardiac development. This approach could confound the analysis of the more evolutionarily diverged anterior neural tube, due to tissue-specific heterochronies (19).

The bulk of our knowledge regarding mammalian telencephalon development stems from studies in mouse. Therefore, it is critical to identify when and where human forebrain development differs from that of mouse. The establishment of fates and relative sizes of the initial telencephalic progenitor areas have been shown to be essential for forebrain development. These early processes are thus likely to play a role in evolutionary divergence of forebrain functions and connectivity. We set out to explore this possibility by comparing telencephalic signaling pathways and their downstream effectors between mouse and human at the crucial patterning stages of development. We hypothesized that, given the differences observed in signaling dynamics of Shh in such closely related species as Lake Malawi cichlid fishes, it would stand to reason that similar or greater differences may exist between human and mouse. To test this hypothesis, we used single-cell RNA sequencing (scRNA-seq) and hybridization chain reaction (HCR) RNA fluorescent *in situ* hybridization techniques to investigate the differences in the expression and downstream effects of the signaling pathways that pattern the telencephalon in human and mouse.

## Results

### Transcriptomic similarities and differences in telencephalon patterning between human and mouse

We performed a detailed comparative study of patterning of human and mouse telencephalon using scRNA-seq. We dissected human and mouse forebrains (removing as much of the diencephalon and retina as possible) from 4 distinct stages for each species (human: CS12, CS13, CS15 and CS17; mouse: E9.5, E10.5, E11.5 and E12.5, Figure 1A). These stages were chosen based on anatomical similarities between human and mouse forebrains and published tables of stage equivalency (20). Each stage contains two separate 10X chromium runs. For mouse, each run corresponds to a sample from a separate individual of the same litter at a given stage. For human, each run corresponds to a technical replicate from the same individual (due to rarity of fresh human embryonic tissue).

After performing quality control and removing low quality cells from each sample separately (figure S1), we integrated all samples together and removed non-forebrain cell types at the first step. This yielded 55,758 cells, of which 27,152 were from human and 28,606 were from mouse. We then reclustered these forebrain cells (Figure 1C) and annotated them, based on common markers (figure S2), into broad identities including 3 signaling populations (ventral, dorsal, and anterior), ventral and dorsal progenitors and neurons. These telencephalic and signaling populations accounted for 42,734 of the cells. Posterior signaling center, progenitors, and neurons (and one unidentifiable cluster), accounted for the remaining cells (Figure 1D).

By comparing the proportion of mouse and human cells in each cluster, we identified a number of progenitor clusters that are human-enriched (figure S3A,D). In particular, two clusters in the early progenitors immediately stand out, one in ventral and the other in dorsal telencephalon progenitors (figure S3B-E). For the cluster among the ventral progenitors, there are some intriguing top differentially expressed genes (DEGs). They are the medial ganglionic eminence (MGE) and septum marker NKX2-1, as well as SFTA3 (NANCI/Sfta3-ps in mouse) and PEG10—although PEG10 expression seems to be widespread in our scRNA-seq data (figure S3B). In human, SFTA3 encodes a 94 amino acid long protein, while, in mouse, there is no open reading frame for this gene (24). We used HCR to probe the expression of SFTA3 in human and Sfta3-ps in mouse to ascertain the difference between the two expression profiles. At CS14, SFTA3 is expressed in the ventral telencephalon and diencephalon, almost mirroring the expression of NKX2-1. The expression pattern of Sfta3-ps at E9.5 closely resembles that of SFTA3 in CS14 (figure S3F). We could not detect any higher expression level of SFTA3 or expansion of its domain in human compared to mouse. The difference obtained in scRNA-seq appears to be due to a lack of 3’ UTR of the mouse non-coding RNA, preventing 10X amplification of the transcript, highlighting the importance of validation experiments to ascertain transcriptomics-based findings.

The dorsally located cluster with a high proportion of human cells has MIR9-1HG/C1orf61 as its highest DEG, as well as EMX2, EMX1, and FEZF2 (figure S3C). FEZF2 is an anterior neural tube specific zinc finger transcriptional inhibitor, which may act downstream of WNT signaling and is upregulated upon overexpression of DKK1 (25). MIR9-1HG RNA has previously been shown to be expressed in the human CS16 cortical progenitors using RNAscope technology (26). Here we detect strong expression of MIR9-1HG transcript in dorsal telencephalon as early as CS13, dorsally co-localizing with EMX2 expression (Figure 3F). This gene has no orthologue in mouse, and we confirm its human expression and identify the cluster in very early human dorsal telencephalon.

### Identifying developmental stage equivalence between mouse and human telencephalon

When comparing the UMAP representations of the scRNA-seq data separated by species (Figure 1E), the striking difference between the two is the cell density of neuronal populations. The mouse samples contribute many more cells to both the ventral and the dorsal neuronal subpopulations than the human, indicating that the assumed embryonic stage equivalence for these species (Figure 1A) does not hold for the telencephalon. To assess equivalence at the local telencephalon level, we compared the eight stages at a transcriptomic level to find where each stage fits within the timeline of telencephalon development. We calculated cosine distances between each stage and all others in the 30-dimensional principal component space of telencephalon progenitors, neurons and signaling centers. We set human CS12 as the earliest stage (the stage at which the forebrain has just completed neural tube closure), in the combined human and mouse series (Figure 2A). This temporal order, based on transcriptomic similarities, deviates from the established order based on body morphological milestones (Figure 2B), indicating a slower developmental tempo of human telencephalon than other embryonic tissues.

**Fig. 2.**
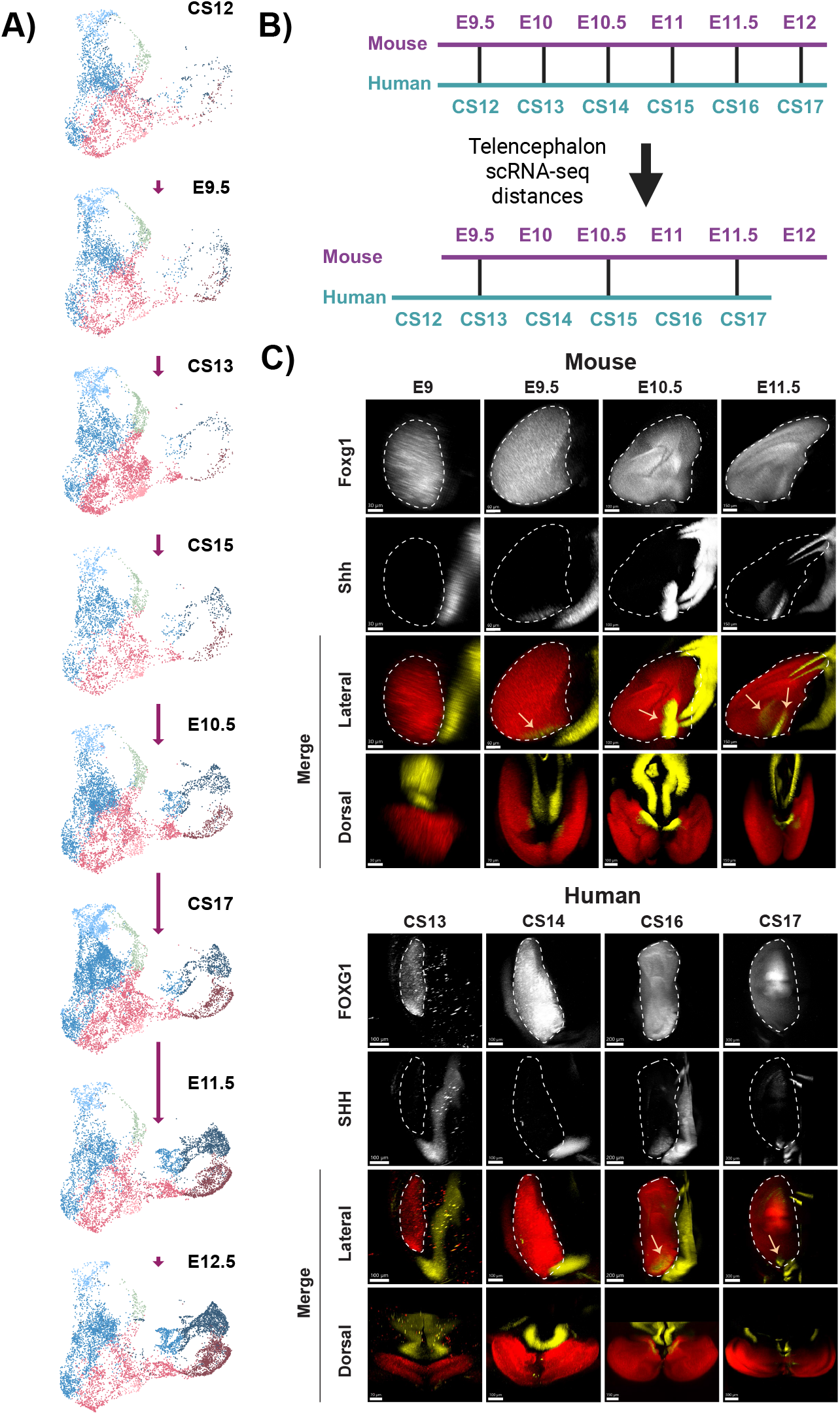
Realignment of human and mouse equivalent stages. **(A)** UMAP plots of integrated human and mouse telencephalon superclasses separated by stage and placed along a timeline from the earliest to the latest. The length of each arrow is scaled to the cosine distance between center of mass of the PCA space of each stage to the next (see Methods for details). **(B)** The equivalency between human and mouse embryonic timelines is realigned to reflect the timeline arrived at in **(A)** for the telencephalon. **(C)** Lateral and dorsal views of mouse and human telencephalon labeled for FOXG1 (red) and SHH (yellow) using wholemount HCR. Four embryonic stages for mouse (E9, E9.5, E10.5, and E11.5) and four for human (CS13, CS14, CS16, and CS17) are shown. Arrows point to the presence and location of SHH expressing regions within the FOXG1 expressing telencephalon. The dashed lines outline the area of FOXG1 expression. Dorsal views are included to clearly indicate non-telencephalic SHH.

To validate the temporal observations made from scRNA-seq analysis, we set out to detect the expression of major determinants of telencephalon patterning. SHH is initially expressed in the hypothalamus ventral to and outside of the FOXG1 expressing telencephalon. As patterning progresses, a discrete ventral telencephalic domain of SHH appears, overlapping with that of FOXG1 (27, 28). We visualized FOXG1 and SHH mRNA expression domains in the two species, using HCR. In mouse, Shh expression is not detected in the telencephalon at E9, but is present at E9.5, with a marked increase in its domain at E10.5. By E11.5, there are two distinct domains of expression within the FOXG1 telencephalon (Figure 2C, top panel). In human, SHH expression is not found in the FOXG1 expressing telencephalon at CS13 and is hardly there at CS14. Its telencephalic expression can only be reliably detected from CS15 onward (Figure 2C, bottom panel). Therefore, wholemount detection of the SHH signaling center demonstrates a delay in the induction of the telencephalic ventral signaling center in human, more pronounced than the delay found in the overall telencephalic transcriptome (Figure 2B).

### Reduced diversity of early progenitor identities in human telencephalon

Having identified the stage equivalence between mouse and human, we asked whether progenitor identities across anteroposterior (AP) and dorsoventral (DV) axes are conserved between the two species at this early stage. Therefore, we examined how progenitor domains are distributed in E9.5 and CS13, by analyzing the scRNA-seq progenitor clusters for each stage separately, then comparing them to each other. Using the same level of resolution for both species, we systematically obtained fewer progenitor clusters in human than mouse. We also noticed an unexpectedly high number of progenitor clusters (12 and 7, respectively) at these early stages. Since we had access to many more mouse E9.5 samples than early human embryonic samples, we focused on validating these clusters *in vivo* at this stage. As scRNAseq clustering was based on an arbitrary— albeit educated—clustering resolution, we compared E9.5 clusters and their DEGs to each other, in pallium and subpallium. We combined mouse clusters expressing very similar known regional markers into a final eight groups (Figure 3A).

**Fig. 3.**
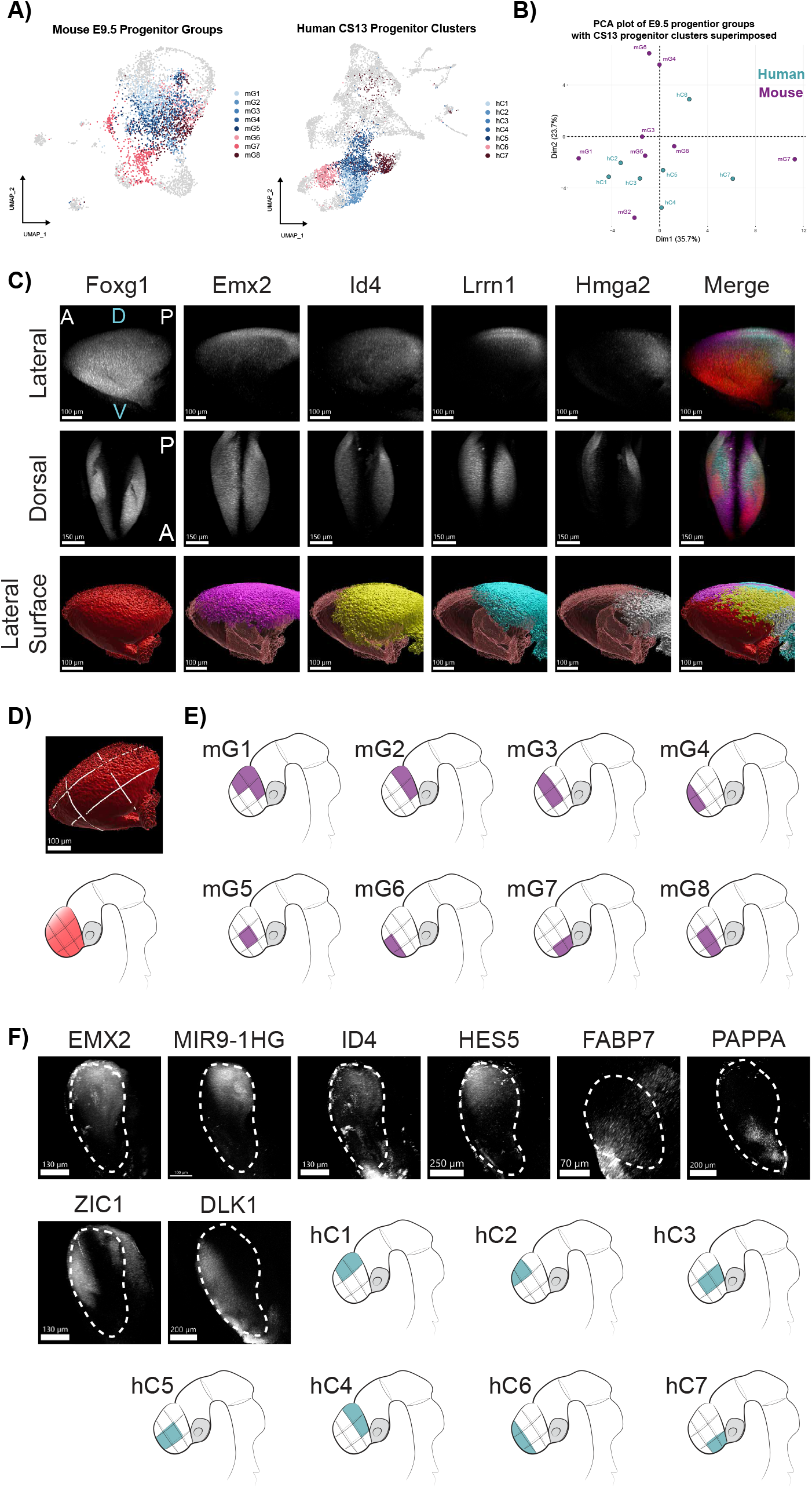
Dorsoventral and anteroposterior distribution of progenitor identities in early mouse and human embryonic telencephalon. **(A)** UMAP plots of mouse E9.5 and human CS13 cells processed separately. Dorsal (shades of blue) and ventral (shades of maroon) progenitor groups (E9.5) and clusters (CS13) are indicated on the UMAP plots. For E9.5, some clusters have been combined into groups (see Methods) and for human single clusters are shown. **(B)** A PCA plot of E9.5 progenitor groups with CS13 clusters superimposed as supplements. First dimension corresponds to the DV axis with dorsal groups to the left and ventral to the right. Second dimension corresponds to the AP axis with anterior groups to the top and posterior to the bottom. **(C)** Lateral and dorsal views of mouse mG1 group telencephalon labeled for Foxg1 (red), Emx2 (magenta), Id4 (yellow), Lrrn1 (cyan), and Hmga2 (gray), using wholemount HCR, and the resultant Imaris surfaces. **(D)** The Foxg1 surface (above) and a schematic of the telencephalon (below) with lines drawn to divide it into 9 zones along the DV and AP axes. **(E)** Schematics of all eight mouse groups (mG1-mG8) and the zones in which their top DEG expressions overlap. **(F)** Lateral views of human EMX2, MIR9-1HG, ID4, HES5, PAPPA, ZIC1, and DLK1 expression in CS13 and FABP7 expression in CS12, as well as schematics showing inferred zones of expression for seven human CS13 progenitor clusters. The dashed lines outline the area of FOXG1 expression.

For each group, the four highly expressed representative genes were selected for validation, in conjunction with Foxg1. In order to visualize all five markers in the same sample, we developed a method (first part of the workflow in figure S4) combining multiplexed HCR staining and a modified version of the immunolabeling-enabled three-dimensional imaging of solvent-cleared organs (iDISCO+) technique (29) for 3D wholemount imaging of embryonic telencephalons. The stained and cleared samples were imaged using a confocal microscope with tunable white-light laser to decrease the cross-talk between fluorophores with overlapping excitation spectra.

We determined the expression profiles of all genes using this workflow. An example for the mouse group mG1 is shown in Figure 3C, and images for the remaining groups are in figure S5A. Expression profiles of all five makers were obtained in the same wholemount sample, and regions of overlap were examined in 3D (Figure 3C). To map the progenitor areas, we divided the telencephalon into 9 zones (Figure 3D), obtained from three DV and three AP divisions. The HCR results validated the groups found by scRNA-seq and placed these onto a 3D map. This mapping reveals the presence, in mouse, of medial identities along the AP axis at this very early stage, unsuspected by previous studies (Figure 3E).

Given the difference in progenitor cluster numbers between E9.5 and CS13, we sought to identify similarities, at the transcriptomic level, aiming to match progenitor populations between the two species. We first generated a principal component analysis (PCA) plot of the E9.5 groups, which represented the DV and AP spread of the eight groups along its first and second components, respectively (Figure 3B). We superimposed the seven clusters from CS13 onto this plot, finding that these clusters spread along the axes in a similar way to the mouse groups. Given the rarity of human samples at these stages, we were only able to validate a small proportion of markers from these clusters to connect them to groups from E9.5 mouse.

The most ventral (hC7), most anterior (hC6), and most posterior (hC4) clusters in CS13 are distinctly in the ventral, anterior and posterior regions of the PCA (Figure 3B). Their top DEGs also align well with the corresponding groups in mouse. The markers found in hC7 (NKX2-1 and MBIP) are present in mG7 and can be seen within the ventral telencephalon as early as CS12 (figure S5B). Anteriorly, hC6 expression of ZIC1 in human (Figure 3F) shows a similar profile to that of E9.5 mG4 (figure S5A). For the most posterior cluster of CS13, hC4, we were not able to validate any main markers, but its top DEG, NR2F1, is shared with the E9.5 mG2.

The other four dorsal clusters of CS13 share a number of top DEGs with each other. The most ventrally situated of these is hC5, highly expressing FABP7, PAPPA, and HES5 (Figure 3F). The CS13 hC3 cluster shares many of the same DEGs with hC5; but, among those it also contains EMX2 and MIR9-1HG which are more dorsally and posteriorly expressed (Figure 3F). The two most dorsal CS13 clusters, hC2 and hC1, have the same dorsal markers, EMX2, MIR9-1HG, ID4, and HES5 (Figure 3F) expressed among their top DEGs. However, when these two clusters are directly compared, hC2 differentially expresses more anterior markers (e.g. ZIC1) compared to more posterior markers (e.g. NR2F1) for hC1. Based on these comparisons, and in conjunction with the validated markers, we generated a map of human telencephalic regions at CS13 (Figure 3F). It points to a similar DV regional diversification, but a poorer fate separation along the AP axis, in human compared to mouse at these early stages of development.

### Human features in dorsal and ventral telencephalic patterning

From these results, it is clear that the human CS13 progenitor DV diversity is mostly dorsal (5 dorsal identities out of 7 clusters) and lacks AP definition. To validate these transcriptomic findings in the 3D brain across developmental time, we measured dynamics of AP and DV patterning using a minimal set of regional markers and signaling molecules. WNT8B is a major signal in the dorsal telencephalon, repressed by FOXG1 (30). EMX2 is induced by WNT8B, expressed in a dorsal-high to ventral-low pattern inside the pallium (31), while PAX6 is initially relatively homogeneously expressed in the prospective pallium, then is inhibited dorsomedially (32). On the ventral side, SHH patterns the subpallium with NKX2-1 as one of its main targets (33). AP patterning is controlled by FGF signaling, with FGF17 being a main player in the rostral patterning center, also defining the anterior-most limit of the telencephalon (34).

Comparative studies using thin tissue sections often lead to difficulties in choosing equivalent sections across species. Therefore, to quantitatively examine, in both species over time, the expression domains of these genes, we followed the same method for 3D wholemount imaging of embryonic telencephalons as above (figure S4). Since fresh human embry-onic samples at the early stages (younger than 8 weeks post-conception) are increasingly scarce, we devised a method to extricate tissue from wax of archival formalin-fixed paraffin-embedded (FFPE) samples that provides a solution to rarity in human tissues across any organ of interest. Additionally, to maximize the amount of data generated, we used each tissue sample for multiple rounds of HCR by stripping old probes with DNase I.

We processed mouse (E9.5-E11.5, n=3 for each stage and each gene) and human (CS12-CS17, see Figure 4B,C for n) embryonic heads, according to this workflow. Images were segmented to compute 3D meshes with the VEDO python module, and we generated custom scripts, based on the Computational Geometry Algorithms Library (CGAL), to compute mesh unions and intersections and calculate their volumes. Figure 4A and video S1 illustrate an example of the resulting images and meshes, showing the first round of HCR imaging of a human CS14 embryonic head stained for five markers. We normalized volume of each marker to the total volume of the telencephalon, defined by the sum of the FOXG1 and WNT8B telencephalic domains (35). Comparison of these territories, between human and mouse, reveals different temporal dynamics.

**Fig. 4.**
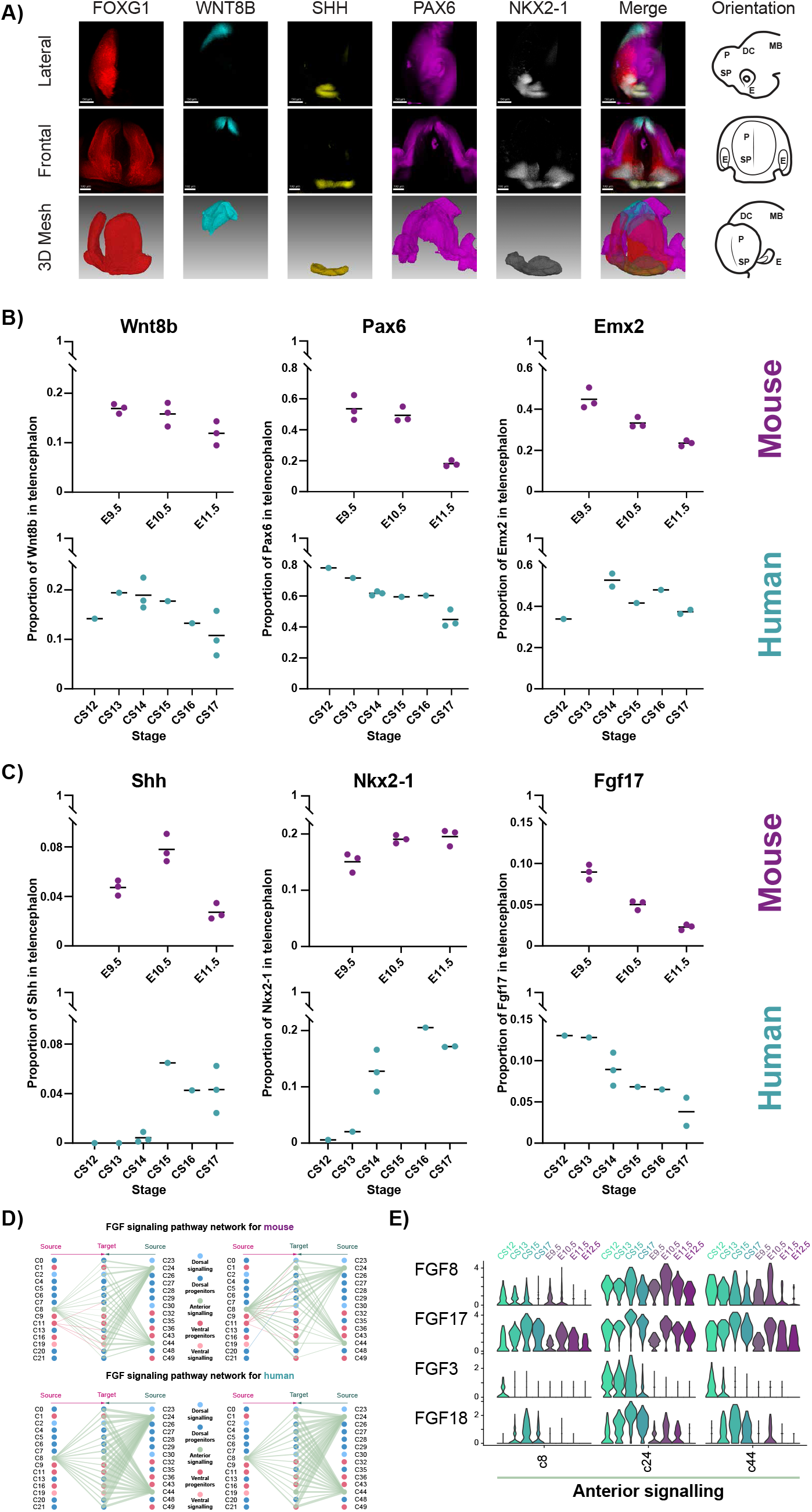
Heterochrony in ventral and dorsal signaling and progenitor development of the human telencephalon. **(A)** Images and meshes for a human CS14 head. The five markers used for HCR are FOXG1 (red), WNT8B (cyan), SHH (yellow), PAX6 (magenta), and NKX2-1 (gray). The right panel shows a schematic of the orientation of the images or meshes. Lateral and frontal views of the HCR stained head and a three-quarter view of the meshes are shown to convey the 3D nature of the images. P: pallium, SP: subpallium, DC: diencephalon, E: eye, MB: midbrain. **(B)** Volumes of dorsal signaling and progenitor markers and **(C)** ventral signaling, ventral progenitor, and anterior signaling markers as a proportion of telencephalon volume. Three mouse (E9.5-E11.5) and six human (CS12-CS17) time points were used. For mouse n=3 samples for each stage were used. For human CS12 n=1, CS13 n=1 (except EMX2, where n = 0), CS14 n=3 (except EMX2, where n=2), CS15 n=1 (except NKX2-1, where n=0), CS16 n=1, and CS17 n=3 (except EMX2, NKX2-1, and FGF17, where n=2). The human data are generated from fresh or archival FFPE samples, and, as such, a number of genes were not detectable for certain samples depending on how long they had been archived or potential damage. **(D)** Comparison of the FGF signaling pathway using CellChat. The plots show the aggregate interaction weights for FGF signaling components for mouse (top) and human (bottom). The thickness of the lines from source clusters to target clusters is proportional to the strength of the interaction. **(E)** Single cell gene expression violin plots for four main ligands in the FGF pathway (FGF8, FGF17, FGF3, and FGF18) showing a comparison of their expression levels over time in anterior signaling clusters in human (teal) and mouse (purple).

The proportion of the dorsal signaling center (WNT8B domain) decreases in a similar fashion from CS13 to CS17 and from E9.5 to E11.5 (Figure 4B). In the pallium, PAX6 expression decreases in mouse over the same period, with a large drop to about 20% of the telencephalon at E11.5 (dorsal to ventral regression with expression kept in ventral pallium). In human, on the other hand, the decrease in size of PAX6 territory is more gradual, dorsally reducing to 40% of the telencephalon at CS17. The difference in dynamics of EMX2 is more pronounced, with a small decrease in proportion in human, to 40% of the telencephalon at CS17, compared to 20% in the dorsal-most pallium at E11.5 in mouse. Therefore, despite a very similar dynamic in dorsal signaling center proportion, the downstream read-out of the signal throughout the telencephalic tissue is different between the two species.

Ventrally, the mouse SHH signaling center is already well within the telencephalon at E9.5, its relative size increasing at E10.5 to then shrink by E11.5 (Figure 4C and Figure 2C). Contrastingly, in human, SHH is not expressed within the telencephalon at CS12 nor CS13, scarcely appearing at CS14. The relative size of the ventral signaling center becomes similar to mouse at the later timepoints. This delay in onset of telencephalic SHH-expressing population is associated to changes in activation of transcriptional targets. Mouse NKX2-1 domain consistently increases from E9.5 to E11.5, while in human, NKX2-1 is almost absent at the earliest stages (CS12 and CS13), finally reaching a similar relative size to the mouse E10.5 domain at the latest CS17 stage. This strongly suggests that the source of SHH secreted from the hypothalamus is not sufficient to establish NKX2-1 expression in the ventral telencephalon at the earliest time points in human. Delay in induction does not seem to have a lasting effect in organization of the ventral progenitors.

Anteriorly, the size of FGF17 expressing anterior neural ridge (ANR, (36)) domain is bigger in human (CS12 and CS13) than mouse at early stages and stays so over time (Figure 4C). However, the pace of decrease of signaling center size is similar between the two species.

Overall, these observations indicate a delay in human ventral signaling center induction and its longer persistence, a similar timing of induction and a bigger size of the FGF-secreting anterior signaling center, and a similar dorsal signaling center size and dynamic. These differences correlate with a species-specific response to signaling inside the pallium, suggesting that this divergence may be due to human-typical spatio-temporal integration of the three signaling centers.

### Identification of a human feature of the ANR signaling center

To deepen our understanding of the human signaling landscape in the nascent telencephalon, we characterized the signaling networks in our scRNA-seq datasets, using CellChat (37), and found additional differences in signaling between mouse and human (figure S6C). We disregarded human only immune-related signaling events, as well as mouse enriched signaling involved in neurogenesis. Confirming our observations so far, this analysis showed that hedgehog signaling is more complex in mouse than human (figure S6A). Focusing on pathways enriched within progenitors and signaling populations, we identified a human signature of non-canonical WNT (ncWNT) and FGF signaling (figure S6B and Figure 4D).

FZD5, is present at higher levels in some human clusters, specifically those annotated as anterior signaling center and ventral progenitors (figure S6D), suggesting an involvement of this receptor in setting up ventral and anterior identities in human. Within the FGF family, while FGF8 expression is similar in the two species, FGF17 is increased in human anterior signaling center clusters. FGF18 and FGF3 expression profiles, however, are very high in human cells within these clusters and barely detectable in mouse (Figure 4E and figure S6D), providing candidate determinants for human features of forebrain patterning.

Having found a pronounced human delay in tempo of ventral signaling (SHH and NKX2-1) and a human ANR signature, we explored their implications for the neurogenic progression of dorsal and ventral cells. We used the transcriptomic approach described in Figure 2A, to find the distances between the dorsal and ventral neuronal populations of each embryonic stage. We excluded the earliest timepoints as they did not contain many neurons, making them more susceptible to noise. For ventral neurons, CS17 is between CS15 and E10.5, while for dorsal neurons CS17 is between E10.5 and E11.5 (figure S7A). Therefore, dorsal neurogenic tempo matches the overall telencephalon timeline, while human ventral neurogenic progression displays a slower tempo (figure S7B). These results indicate that human signaling characteristics and delay in specification of ventral progenitor domains leads to slower progenitor maturation and ventral neurogenesis.

## Discussion

In this work, we have taken advantage of scRNA-seq and 3-dimensional HCR techniques to identify major signaling pathways determining the early patterning of human telencephalon. We have shown that the development of human telencephalon is slower than the rest of the embryo, in particular those organs used for determining stage equivalencies with mouse. The delay in telencephalon development we found using our scRNA-seq dataset (Figure 2B) is in agreement to that found in the human spinal cord by Rayon and colleagues who used similarity in sizes of areas of well-known marker genes to arrive at their equivalency (38). Likewise, telencephalon development appears to be further delayed in ferret when compared to mouse (39), hinting at a common mechanism in development of complex neural tissue.

The human telencephalon development, however, not only displays heterochrony compared to other embryonic structures, but also shows developmental heterochrony along its DV and AP axes when compared to mouse equivalent telencephalic stages.

We leveraged the transcriptomic data from our scRNA-seq datasets to probe the diversity of human and mouse progenitors. In mouse E9.5, we had annotated five of the eight groups as dorsal with the other three as ventral based on their top DEGs. Our validation showed that, ventrally, laterally, and dorsally, the groups were spread along the AP axis (Figure 3E, Figure 5A). This early separation of populations along the AP axis of the telencephalon had not been readily detectable in the transcriptomic data, potentially due to a lack of extensive characterization of AP patterning markers at such an early stage. While efforts are being made in this direction, they are still mostly focused on later stages (40). Comparing clusters from human CS13 to those validated in the mouse equivalent stage pointed to a similar DV diversity, while anteroposteriorly, the clusters are less defined in human (Figure 3F, Figure 5A), showing lack of fate definition of medial AP identities.

**Fig. 5.**
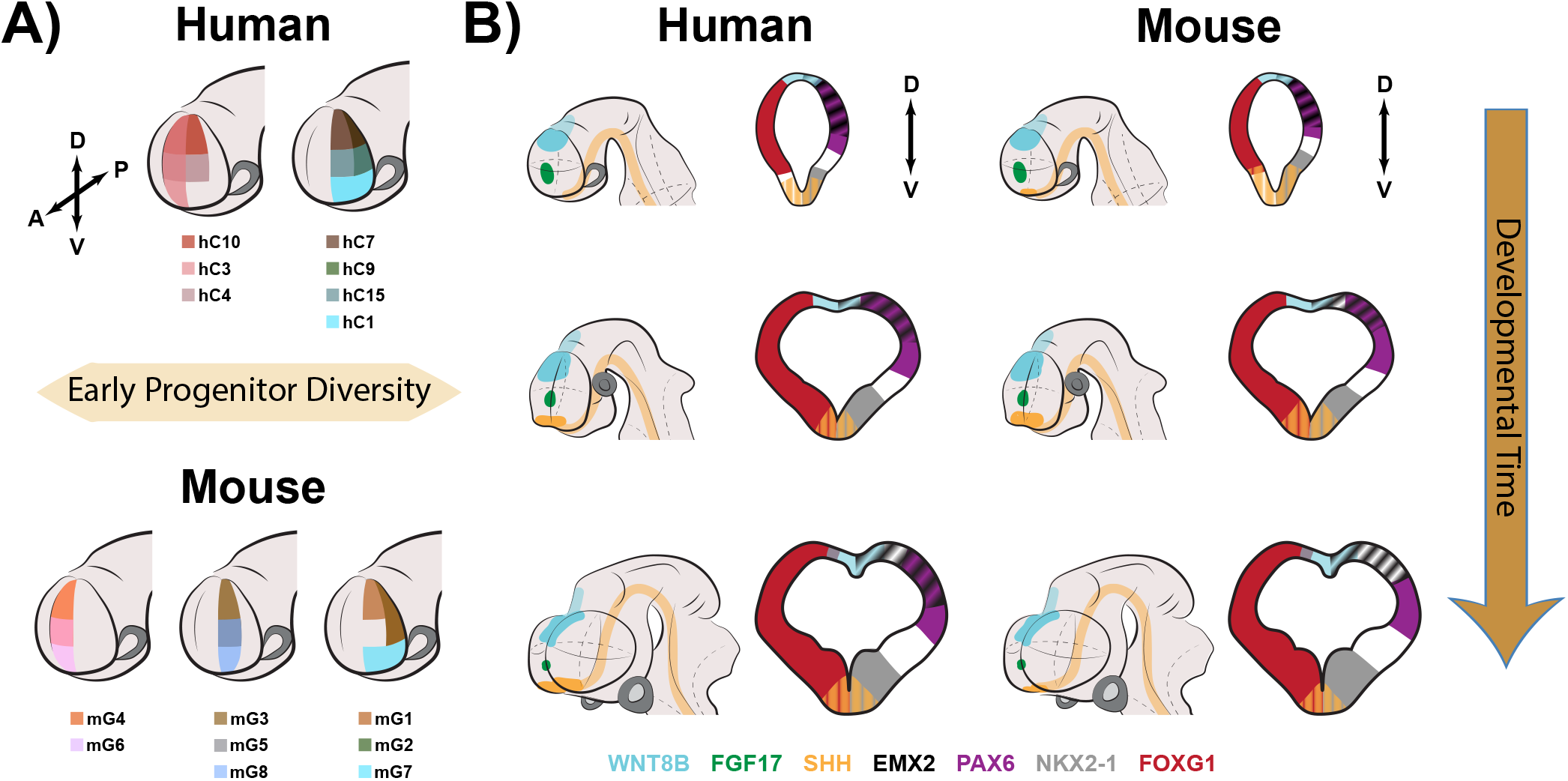
A schematic summarizing early differences in human and mouse telencephalic signaling and progenitor diversification. **(A)** Early during the phylotypic period, mouse telencephalon progenitors are spread along the DV and AP axes in transcriptomically distinct populations. In human, a similar spread is seen dorsoventrally, but to a lesser extent anteroposteriorly. **(B)** These differences coincide with a larger volume of dorsal telencephalic regions in human and a delay in ventral signaling, underpinning a correlated heterochrony along the DV and AP axes of human telencephalon when compared to mouse.

We further investigated the establishment of DV patterning over time and showed that the human telencephalon retains a larger dorsal identity domain for longer (Figure 5B). This prolonged maintenance of the dorsal identity domain does not appear to be correlated with a modified dorsal signaling by WNT8B. Instead, it matches a delay in the induction of the NKX2-1-expressing region and establishment of SHH signal in the ventral telencephalon (Figure 4B,C). This delay may provide the means to establish a wider set of dorsal identities inside the pallium later.

The delay in ventralization of the telencephalon does not lead to a large difference in the proportion of ventral (subpallial) and dorsal (pallial) regions at the end of the period studied. The volume of the NKX2-1 expressing region catches up with that of mouse by the final timepoints. The delayed ventralization of the telencephalon in human raises questions about how this may affect diversification of progenitor identities at the earliest stages of development. As it is correlated to the slower establishment of, and differences in, the human progenitor identities along the AP axis, it points to a role for the ventral SHH signal in integration of DV and AP patterning during telencephalon development.

Taking advantage of the CellChat tool we identified other signaling pathways contributing to the human telencephalic signaling signature (figure S6). We specifically focused our analysis on those pathways whose components were differentially present in ventral or anterior progenitor and signaling clusters, as these would be most likely to interact with SHH and/or its downstream effectors. While FGF8 and FGF17 expression levels, at the ANR (12, 41, 42), are similar across species, FGF3 and FGF18 are highly expressed in human cells from the earliest stage, and are only detectable at much lower levels in mouse in these same clusters. FGF18 expression has not been described in human at early stages of development (43), our work providing the first evidence for an early involvement in human telencephalic features. This ligand is shown to inhibit Shh expression in mouse embryonic palatal shelf (44). In mouse, Fgf3 does not seem to have a role in early forebrain development (45), while the expression of Fgf18 overlaps that of Fgf8 rostroventrally inside the ANR (46). The abundance of FGF3 and FGF18 expression differentially found in human anterior clusters (Figure 4E), therefore hints to a role for them in delaying SHH induction, its subsequent ventral spread, and activation of its effectors in human. Suitable *in vitro* human models of telencephalon DV patterning, such as those being currently developed ((47) and unpublished data) will be needed to test this hypothesis.

As intriguing is the differentially enriched expression of FZD5 in human ventral-most progenitor and ANR clusters and in two of the lateral dorsal clusters. Fzd5 is expressed in mouse from at least E9.5 in ventral telencephalon, continuing its expression beyond the timepoints we have examined (48). To our knowledge, no interaction between FZD5 and SHH are reported to date, while Fzd5 expression is negatively regulated by Nkx2-1 in mouse ventral telencephalon (49), pointing to a potentially complex regulatory relationship between ncWNT and SHH signaling.

In conclusion, we have found that ventral SHH signaling is profoundly delayed in human telencephalon, while FGF3 and FGF18 are uniquely expressed in the human anterior signaling center. These distinct signaling features are correlated to a reduced AP patterning resolution and delay in ventral neurogenesis in human. The delayed ventralization of the human telencephalon does not lead to a decrease in proportion of MGE within the telencephalon by the end of CS17, but delays its maturation, which is consistent with studies showing a prolonged interneuron generation in human MGE (50, 51). The functional relationship between these human features will be deciphered in future refined *in vitro* telencephalic 3D models.

## Materials and Methods

### Human embryonic tissue

Fresh or formalin-fixed paraffin embedded (FFPE) human embryonic heads were obtained from the MRC/Wellcome Human Developmental Biology Resource (HDBR; https://www.hdbr.org), who gained UK ethics committee approval and written consent from donors. Fresh tissue samples were collected from HDBR at UCL Institute for Child Health and FFPE samples were collected from HDBR Newcastle University Institute of Human Genetics. Embryos were staged based on the Carnegie Stage (CS) system and karyotyped by HDBR staff. Fresh tissue samples were kept in Hibernate-E medium (A12486, Gibco) supplemented with 2% B-27 (17504001, ThermoFisher Scientific) and 2.5 mL/L Glutamax (3505006, Gibco) on ice until processed for scRNA-seq or fixed for imaging. Samples destined for imaging were fixed in 4% paraformaldehyde (PFA) for 1-2 hrs at room temperature depending on size and immediately processed for HCR.

### Mouse embryonic tissue

All animal procedures were carried out at the Francis Crick Institute in accordance with the Animal (Scientific Procedures) Act 1986 under the Home Office project license PPL PP8246537. CD-1 wildtype mouse embryos were obtained at different stages and kept on ice and processed similar to human samples.

### Telencephalon dissection and cell dissociation

Fresh human (CS12, CS13, CS15, CS17) and mouse (E9.5, E10.5, E11.5, E12.5) heads were kept on ice until dissections. The telencephalon was dissected in Hanks Balanced Salt Solution without calcium and magnesium (HBSS, 14185045, Life Technologies) supplemented with 5% heat-inactivated fetal bovine serum (FBS), with an angled micro knife (Fine Science Tools, 10056-12) using the optic vesicle (or eye depending on stage) and thalamic landmarks for guidance. As much of the olfactory epithelium as possible was removed. In all cases, keeping all of the telencephalon intact was prioritized over removing the surrounding tissue.

The telencephalic tissue was dissociated using the same method previously described for spinal cord (52). Briefly, dissected samples were chopped into smaller pieces (depending on size) and incubated in FACSmax (Amsbio, T200100) cell dissociation solution containing up to 300 U/mL Papain (Roche, 10108014001) at 37 ^*°*^C for 10-15 min, depending on size of pieces. After removal of dissection solution, an HBSS solution with 5% FBS, 10 *µ*M Rock inhibitor (Stemcell Technologies, Y-27632), and 1X non-essential amino acids (ThermoFisher Scientific, 11140035) was added, and cells were disaggregated by repeated pipetting (up to 20 times). Single cells were sequentially filtered through 35 *µ*m (Corning, 352235) and 20 *µ*m (Sysmex, 04-004-2325) mesh strainers to remove clumps. Cell quality and viability were assessed using an EVE cell counter (NanoEntek), and only samples with >80% viability were processed for scRNA-seq.

### Single cell RNA sequencing

Single cell suspensions from each mouse and human sample were loaded into one and two lanes, respectively, of a Chromium Chip G (10X Genomics, PN-1000120) of 10X Chromium Controller (PN-120270). The remaining process was carried out according to the manufacturer’s instructions using a Chromium Next GEM Single Cell 3’ GEM, Library & Gel Bead Kit v3.1 (PN-1000121). The resulting cDNA was sequenced on the HiSeq4000 system (Illumina).

### Read preprocessing and alignment

Short reads stored in FASTQ file formats were pre-processed and aligned to custom-built transcriptomes using the CellRanger software (v3.0.2 or v3.1.0). These transcriptomes included an additional chromosome feature containing the GFP-tdTomato sequence (GRCh38-3.0.0_GFP-tdT or mm10-3.0.0_GFP-tdT). The gene expression counts from predicted cells were stored in compressed H5 format (filtered_feature_bc_matrix.h5) and were subsequently imported into R for downstream analysis.

### Single cell gene expression analysis

The analysis of single-cell gene expression data was done primarily in R (v4.2.0) using the Seurat package (v4.1.1) (53). Gene expression data from each experiment were individually converted into an R sparse matrix using the *Read10X_h5* function. Gene feature names from mouse experiments were converted into homologous human gene names to facilitate downstream data integration and comparative analysis. The expression counts from duplicated humanized genes, due to mouse gene duplications, were summed and expression counts from non-homologous genes were removed. The resulting count matrices were used as inputs to create Seurat objects that facilitate the storage and processing of single-cell gene expression data.

Single cells stored in the Seurat object were QC’ed for total genes detected, the presence of doublets (using *scDblFinder* package), the proportion of mitochondrial genes expressed and the annotation of cell cycle phases. Cells with gene counts fewer than 2 median absolute deviation from the population gene count, and with more mitochondrial content than 10% were removed. A second round of cell filtering was done to remove poor quality cell clusters. Essentially, gene counts were normalized using log1p(CPM) method and briefly clustered using the graph-based Louvain method at a resolution of 1.8. Clusters with gene counts below 2000 or RNA counts below 4000 were removed. The expression counts from the remaining high quality cells were normalized using the SCTransform (v0.3.3) method which includes the regression of cell cycle variabilities (from S.Score and G2M.Score metrics).

To determine the identity of the cells and to retain neural cells from the forebrain, we briefly clustered the single cells from each experiment using a resolution of 1 and computed the top differentially expressing genes from each cluster using the *FindAllMarkers* function. We manually curated the cell annotation of each cluster by comparing its top gene markers to well-known markers of various cells in the developing brain. Non-neuronal clusters were subsequently removed from individual Seurat objects.

To unify the mouse and human cell populations from the various developmental timepoints, we performed data integration using functions within the Seurat package. Essentially, the top 2000 overlapping gene features were extracted using the *SelectIntegrationFeatures* function, followed by a recalculation of SCTransform residuals of selected integration gene features using the *PrepSCTIntegration* function. Transcriptomically-similar cells from the various experiments were then mapped using the *FindIntegrationAnchors* function, and a unified, batch-corrected matrix of expression values was obtained using the *IntegrateData* function. To project this integrated single-cell data, we performed PCA followed by UMAP dimensional reduction techniques. Cell clustering using the Louvain method was then performed using PCA embeddings as input and iterated over a range of resolutions (0.4 to 3.0). The best clustering resolution was selected by visualizing the changes in clusters using the *clus-tree* package.

To refine the identities of each cell cluster, we obtained the top differentially expressing genes from each cluster using the *FindAllMarkers* function. We then ranked cluster-specific differentially expressing genes based on both their average log2 fold change and difference in percentage of a gene expressing in that cluster compared to all others (i.e. *avg*_*log*2*FC*\* (*pct*.1*− pct*.2)). We then used the top genes with distinct regional or differentiation stage expression patterns to assign one of 10 broad identities—Ventral signaling, Ventral Progenitors, Ventral Neurons, Anterior signaling, Dorsal signaling, Dorsal Progenitors, Dorsal Neurons, Posterior signaling, Posterior Progenitors, Posterior Neurons, and Non-forebrain/Ambiguous.

### Calculating transcriptomic distances between stages

To find distances between stages and generate temporal timelines, we removed posterior and non-forebrain/ambiguous cells. We then carried out PCA on the remaining telencephalic and signaling cells, and reclustered them. We checked the cluster identities again to ensure that labels had transferred smoothly, which they had, except in the case of one cluster that was non-telencephalic and was removed from further analysis. Based on the mean of the first 30 PCs of the integrated data, we calculated pairwise cosine distances between each stage and all others. We placed CS12 as the first stage, and in a stepwise fashion moved to the next closest stage based on calculated cosine distances. We repeated this process until we reached E12.5 as the final stage. Arrows in Figure 2A represent the cosine distance from each stage to next normalized to the longest distance. We performed the same analysis for figure S7A, using only ventral or dorsal neurons separately, setting the start point at CS15.

### Removal of tissue from paraffin wax

Paraffin was dissolved by incubating FFPE samples in xylene at 60 ^*°*^C twice for 1 hr each. The samples were washed twice more in xylene at room temperature for 1 hr each time. Xylene was removed by washing the samples in 100% ethanol three times 10 min at room temperature, followed by overnight incubation in ethanol. These samples entered the HCR process at the stage before rehydration.

### Wholemount HCR and tissue clearing

Probe sets for each gene were designed using Pybridizer (https://github.com/ctucl/pybridizer, (54)) and ordered as custom pooled oligonucleotides (Integrated DNA Technologies). HCR v3.0 hairpin pairs (B1-546, B2-514, B3-488, B4-594, B5-647) and buffers (hybridization, wash, and amplification) were obtained from Molecular Instruments (MI). Samples were processed based on MI HCR v3.0 protocol for wholemount mouse embryos with some modifications. In brief, after fixing samples were dehydrated in graded series of increasing methanol concentration. Samples were kept in 100% methanol at -20 ^*°*^C overnight or until use. Samples (including those dewaxed) were gradually rehydrated in phosphate-buffered saline with 0.1% Tween-20 (PBST). Samples were permeabilized in 10 *µ*g/mL proteinase K (for 8-20 min depending on size) and post-fixed in 4% PFA for 20 min at room temperature.

Hybridization was carried out overnight at 37 ^*°*^C in 500 *µ*L hybridization buffer, containing 10-40 pmol (depending on number of probe pairs in a set) probes for up to 5 target RNAs. Samples were washed in wash buffer, and then incubated overnight at room temperature in 500 *µ*L amplification buffer containing 30 pmol of each hairpin, snap-cooled immediately before addition to the buffer. Samples were washed in 5X saline-sodium citrate buffer with 0.1% Tween-20.

Samples were embedded in 1% low-melting point agarose and dehydrated in increasing concentrations of methanol or ethanol. They were then delipidated in a mixture of 66% dichloromethane (Sigma-Aldrich, 270997) and 34% methanol for at least 3 hr (up to overnight) at room temperature. The samples were washed twice for 15 min in dichloromethane, followed by refractory index matching in ethyl cinnamate (Sigma-Aldrich, 112372) (55). All samples were kept in ethyl cinnamate in dark at room temperature until imaging.

For samples undergoing multiple rounds of imaging, ethyl cinnamate was removed and samples were washed twice in 100% ethanol and rehydrated in PBST. Probes and hairpins were removed by overnight incubation in 0.2 U/*µ*L DNase I (ThermoFisher Scientific, 4716728001). Samples were washed four times (1 hr each) in PBST, and then processed again starting from the amplification step of the HCR protocol. Samples were processed for a maximum of three rounds of HCR, clearing and imaging.

### Microscopy and linear unmixing

All samples were imaged using a Leica SP8 confocal microscope, equipped with a tunable White Light Laser (Leica Microsystems). The following illumination wavelengths and detection windows were used for each fluorophore. Alexa-488: Ex 498 nm, Em 500-533 nm; Alexa-514: Ex 519 nm, Em 533-561 nm; Alexa-546: Ex 561, Em 569-602; Alex-594: Ex 591, Em 615-657; Alexa-647: Ex 650, Em 658-720. These excitation and emission wavelengths were used to maximize detection while minimizing overlap of signal between nearby channels. Nonetheless, due to large number of channels in a relatively small spectrum, fluorescent signal bled through from each channel into its neighbors. To correct for this bleed through, we performed wholemount HCR and clearing for one gene at a time in E10.5 mouse telencephalons. We imaged these in all channels to find the amount of bleed through for each fluorophore. We then used the linear unmixing analysis module of the LAS X software (Leica Microsystems) to remove overlapping signals and used corrected images for all downstream analysis.

### Segmentation, 3D mesh generation, and marker volume proportion calculation

To measure the volumes of regions expressing a certain marker gene, we first used the ILASTIK software (56) to segment areas of expression in each channel for each stage. We imported the segmentation files for all markers of one sample into a custom-written python script for further processing. The segmentations were binarized and used to generate meshes with the VEDO python module (https://vedo.embl.es). We defined the telencephalon as the volume of FOXG1 and WNT8B. As WNT8B is also expressed outside the telencephalon, when needed, we cut its mesh in such a way to align with the most posterior border of FOXG1.

We generated a custom-written C++ program based on CGAL (https://www.cgal.org) to find mesh unions or intersections. The telencephalon mesh was created by generating a union of FOXG1 and WNT8B meshes. For markers that were imaged in round 1 (FOXG1, PAX6, SHH, WNT8B, and usually NKX2-1), the marker mesh was intersected with the telencephalon mesh to find its volume within the telencephalon. As WNT8B was only imaged in round 1, meshes for other markers (EMX2, FGF17, and occasionally NKX2-1), imaged in other rounds, were intersected with FOXG1 meshes of the corresponding round. As FOXG1 volume would be the same in every round, we used the amount of change in FOXG1 volume from round 1 to other rounds to calculate the volume of telencephalon at other rounds. Essentially, *V*_TCn_ = *V*_TC1_* (*V*_Fn_*/V*_F1_), where *V*_TCn_ is volume of telencephalon in the n^th^ round, *V*_TC1_ is volume of telencephalon in the first round, *V*_Fn_ is volume of FOXG1 in the n^th^ round, and *V*_*F*1_ is volume of FOXG1 in the first round. Finally, we calculated the proportion of each marker within the telencephalon by dividing the volume of the intersected mesh for that marker by the volume of the telencephalon for round the corresponding to that marker.

### Determination of progenitor groups and validation

Fore-brain cells from E9.5 and CS13 stages were subset and clustered separately using the same resolution. Clusters were annotated as described above. For the E9.5 stage, each ventral or dorsal progenitor cluster was compared to each cluster within its region (ventral or dorsal, respectively) in a pairwise fashion, using Seurat *FindMarkers* function, to identify DEGs between clusters of the same region. Those clusters whose pairwise DEGs did not include regionally expressed genes were combined into groups, yielding three and five groups in the ventral and dorsal annotated progenitor clusters, respectively. The top most DEGs for each group were the common top most DEGs in their constituent cluster(s). We chose the top four DEGs for validation, plus Foxg1, using wholemount HCR. Two probe sets (for Mest and Chic2) did not work, so groups mG4 and mG8 had three genes for validation.

After imaging each sample and linear unmixing (as described above), Imaris software (Oxford Instruments) was used to readily generate surfaces for each channel. The surface generated in the Foxg1 channel was subdivided into nine zones by finding the anteroposterior and dorsoventral midlines and dividing each into three. Then the region of overlap for all validation genes within the Foxg1 surface was determined in 3D within the Imaris software and matched to one or more zones.

### PCA of E9.5 groups and CS13 clusters

A common set of genes between E9.5 and CS13 was generated, by finding the intersection of the most highly variable genes from each Seurat object, and their expression in each group or cluster was extracted from the E9.5 and CS13 Seurat objects. The first 30 PCs based on their expression was calculated for E9.5. The coordinates of the CS13 clusters within the E9.5 PC space were predicted as supplementary groups, using the PCA function of FactoMineR package in R. For ease of readability, CS13 cluster numbers were changed from the original Seurat cluster IDs in the following way: hC1 = cluster 7; hC2 = cluster 10; hC3 = cluster 15; hC4 = cluster 9; hC5 = cluster 4; hC6 = cluster 3; hC7 = cluster 1.

### CellChat *analysis*

We processed the data according to authors’ instructions (37) and available tutorials. The human and mouse cells were separated in the integrated dataset and were used to generate two CellChat objects with the same cluster identities. These CellChat objects were then subset to only include progenitor and signaling clusters. The identified information flow of each significant signaling pathway was compared between the two species (figure S6A) to select pathways of interest for further examination using gene expression violin plots, separated by species, for each cluster.

## ACKNOWLEDGEMENTS

We are grateful to Oscar Marin, James Briscoe, Teresa Rayon, and members of the Human Developmental Biology Initiative (HDBI) consortium for invaluable insights and constructive feedback about this work. We thank Alain Chedotal and his group for support in establishing the tissue clearing and 3D imaging methodology. We are grateful to the Human Developmental Biology Resource staff for their tireless efforts to make human embryonic samples available to us. We acknowledge the crucial support of Genomics STP, Advanced Light Microscopy STP, and the Biological Research Facility at the Francis Crick Institute. Parts of Figures 1,2, and 4 were created in BioRender.

This research was funded by: Wellcome Trust Investigator Award WT 220861/Z/20/Z; Human Developmental Biology Initiative (Wellcome Trust, 215116/Z/18/Z); The Francis Crick Institute, which receives funding from Cancer Research UK (FC0010089), Medical Research Council (FC0010089), and the Wellcome Trust (FC0010089); BBSRC LIDo studentship grant number BB/T008709/1; and EMBO Postdoctoral Fellowship ALTF 1053-2022.

## Supplementary Figures

**Fig. S1.**
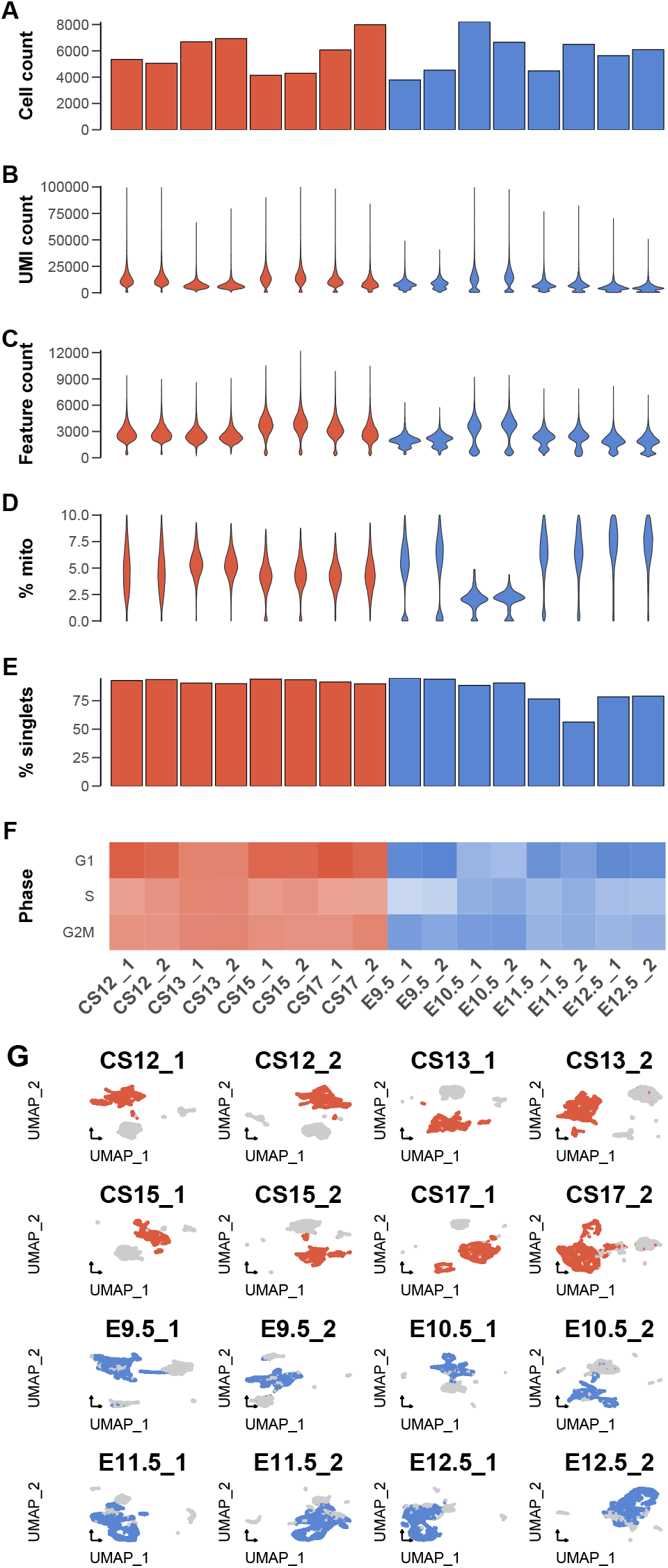
Quality control metrics of cell populations from single-cell RNA-sequencing experiments. **(A-F)** The quality statistics of mouse (blue) and human (red) cell populations from each single-cell experiment. These statistics include cell count **(A)**, RNA UMI count **(B)**, gene count **(C)**, proportion of mitochondrial RNA **(D)**, proportion of singlet cells **(E)** and cell cycle phases **(F). (G)** UMAP embeddings of all cells analyzed from each experiment. Data points colored in red or blue were annotated as neural cells and were kept throughout subsequent analyses.

**Fig. S2.**
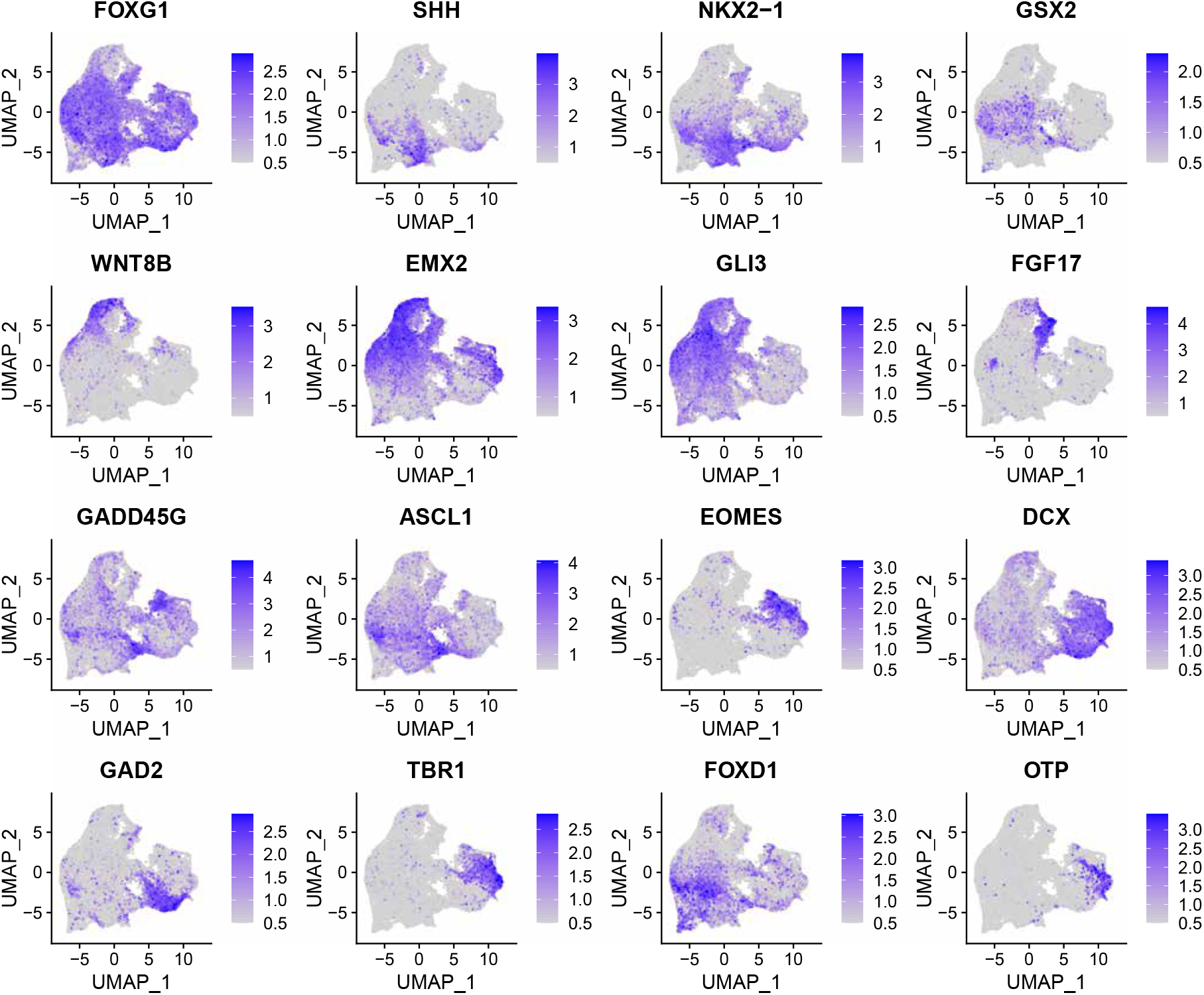
UMAP plots of some major markers used to annotate scRNA-seq clusters of the integrated human and mouse forebrain cells. FOXG1 marks the telencephalon, SHH ventral signaling, NKX2-1 ventral MGE progenitors and neurons, GSX2 ventral LGE progenitors, WNT8B dorsal signaling, EMX2 dorsal medial progenitors, GLI3 dorsal and lateral progenitors, FGF17 anterior signaling, GADD45G intermediate progenitors, ASCL1 ventral intermediate progenitors, EOMES dorsal intermediate progenitors and early neurons, DCX neurons, GAD2 ventral neurons, TBR1 dorsal neurons, FOXD1 diencephalon, OTP hypothalamic neurons.

**Fig. S3.**
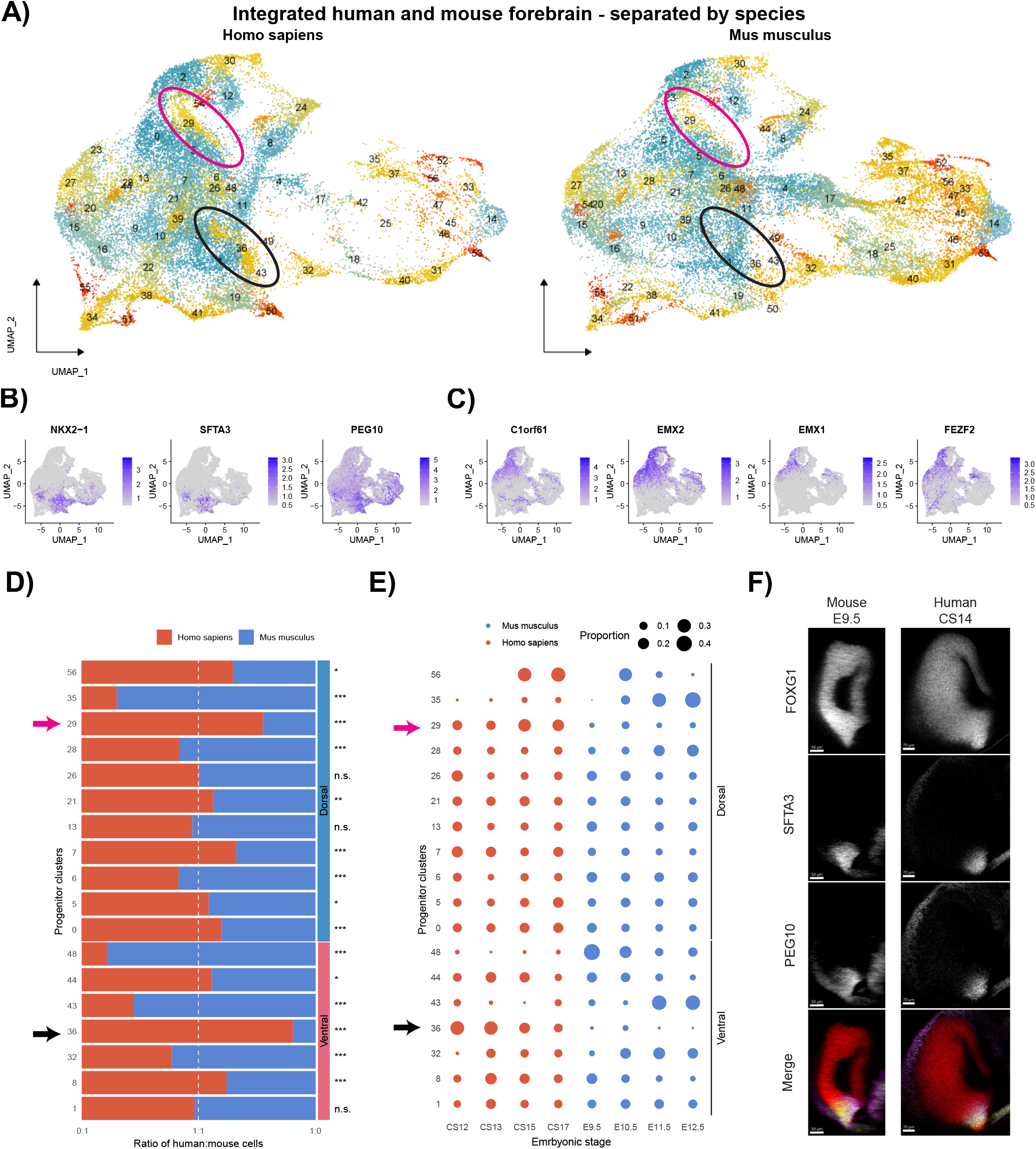
Differences between human and mouse progenitor cell clusters. **(A)** UMAP plots of integrated human and mouse forebrain clusters separated by species. Two clusters (29 and 36) within the dorsal and ventral progenitor populations are indicated with magenta and black ovals, respectively. **(B)** UMAP plots showing three of the top DEGs for the ventral progenitor 36. **(C)** UMAP plots showing four of the top DEGs for the dorsal progenitor 29. **(D)** A stacked bar plot of the ratio of human (red) and mouse (blue) cells in each of progenitor clusters from the integrated dataset. The two clusters enriched for human cells are indicated with arrows. **(E)** A dot plot showing number of cells contributed by each human (red) and mouse (blue) embryonic stage to the progenitor clusters. The two clusters enriched for human cells are indicated with arrows. **(F)** Lateral views of mouse E9.5 and human CS14 telencephalons labeled for FOXG1 (red), SFTA3 (yellow), and PEG10 (magenta) using wholemount HCR.

**Fig. S4.**
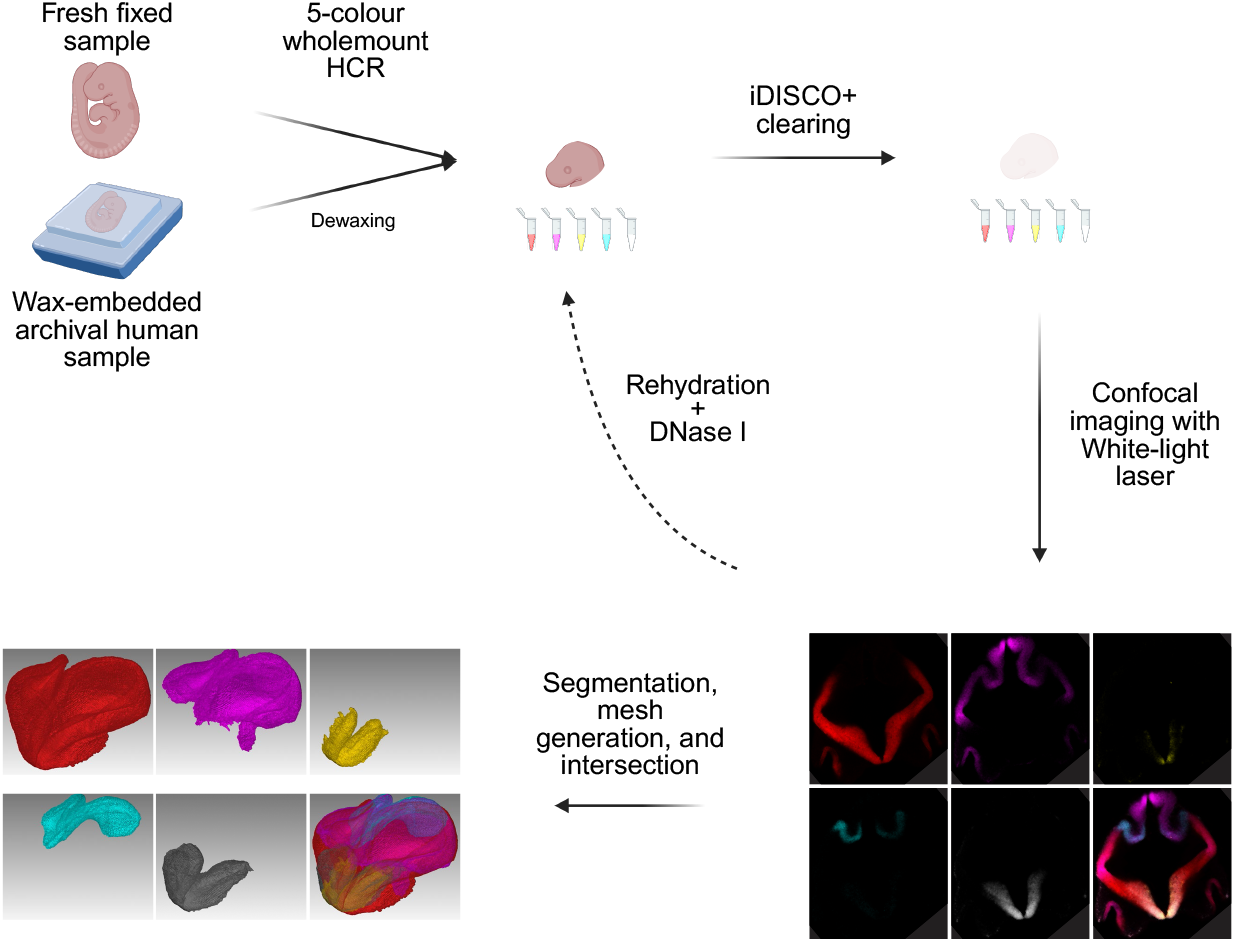
A workflow for wholemount imaging of embryonic telencephalons and measuring regional volumes. A schematic of the optimized workflow from fresh or FFPE archival samples to 3D meshes. This workflow includes dewaxing for FFPE samples, 5-color wholemount HCR staining, iDISCO+-based tissue clearing, confocal imaging, post-processing, mesh generation, and mesh intersection.

**Fig. S5.**
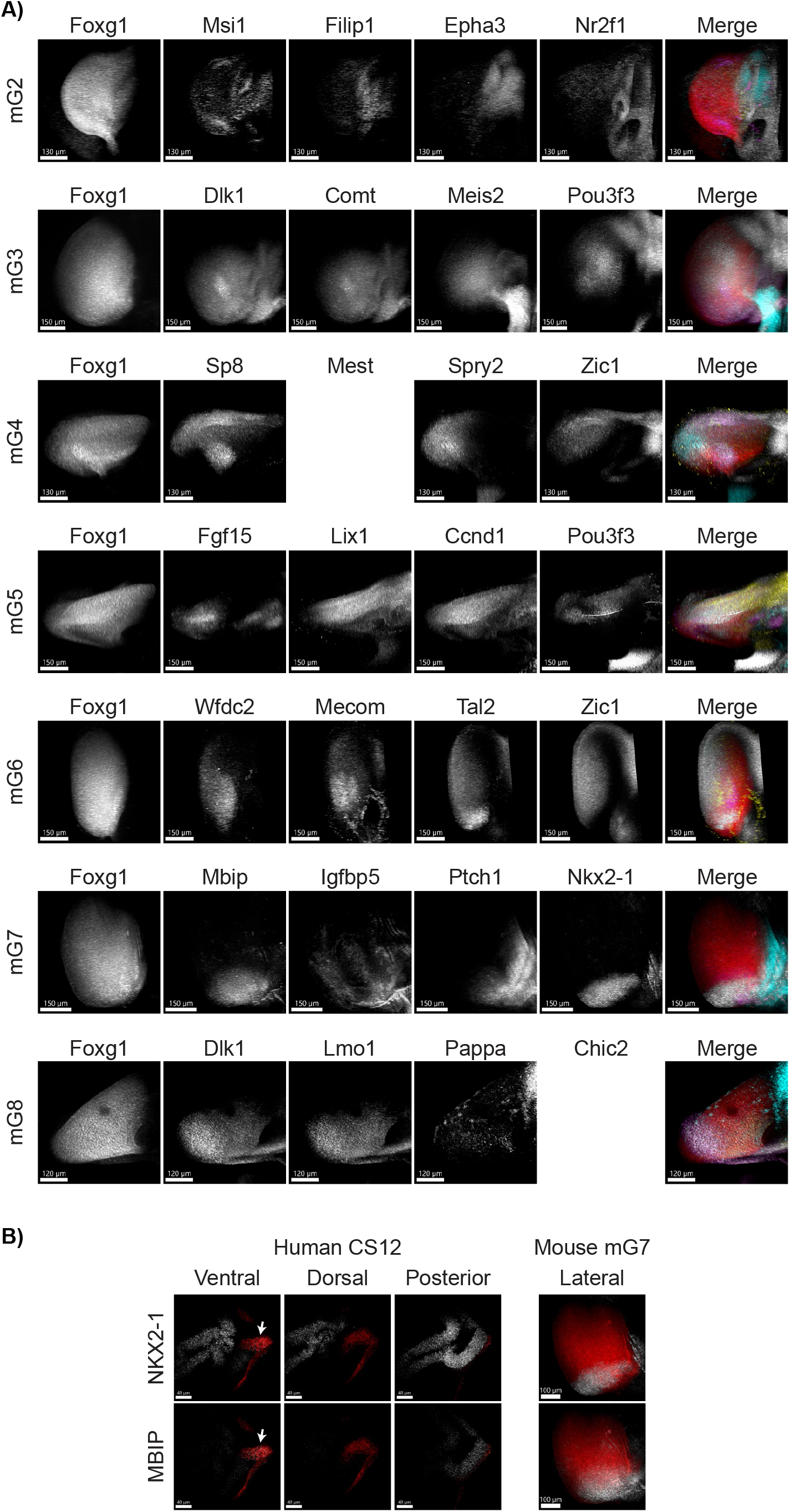
HCR images of progenitor populations in mouse and human. **(A)** Lateral views of mouse mG2-mG8 telencephalons labeled for Foxg1 (red), and four top DEGs in that group. In mG4 and mG8 probe sets for Mest and Chic2, respectively, did not work and are left empty. In the merged column for each group, genes in the second to fifth column are labeled in magenta, yellow, cyan, and gray, respectively. **(B)** Overlap in expression of NKX2-1 and MBIP (gray) within FOXG1 (red) in ventral telencephalon of human CS12 (left) and mouse E9.5 (right) is shown by the white arrow. There is no overlap in dorsally expressing FOXG1 or posteriorly expressing NKX2-1 and MBIP. Note that mouse images are the same as those in **(A)** mG7, shown here again without other markers for clearer comparison.

**Fig. S6.**
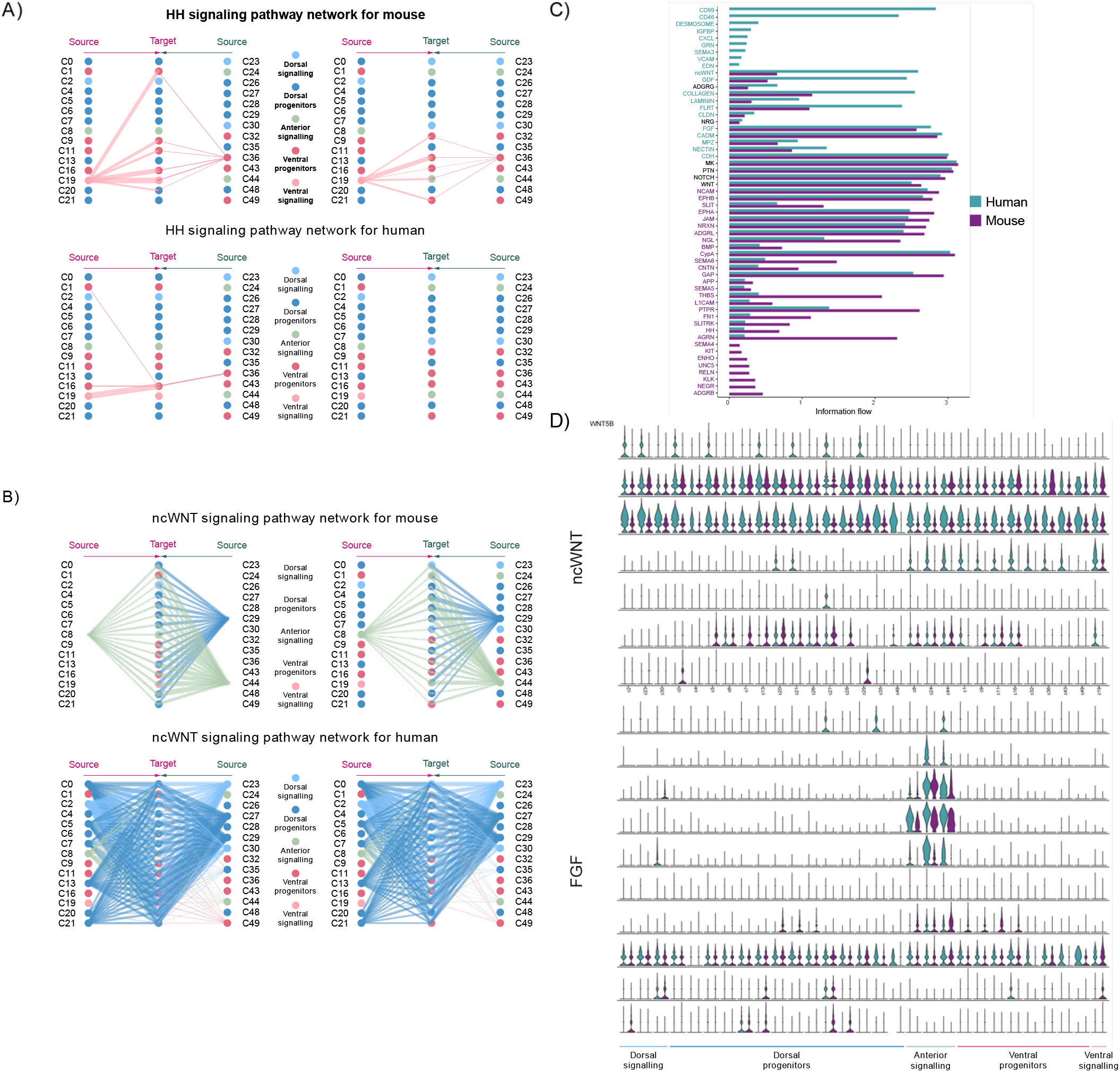
CellChat analysis for comparison of signaling communications in human and mouse. Comparison of hedgehog (HH) signaling **(A)** and non-canonical WNT (ncWNT) signaling **(B)** pathways using CellChat. The plots show a comparison of interaction weights of HH and ncWNT signaling for mouse (top) and human (bottom). The thickness of the lines from source clusters to target clusters is proportional to the strength of the interaction. **(C)** Ranking of all significant pathways based on their overall information flow. The pathways are ordered from stronger interactions in human (teal) to stronger interactions in mouse (purple). **(D)** Single cell gene expression violin plots for ligands and receptors of ncWNT (top) and FGF (bottom) pathways that demonstrate significant interactions in either human or mouse. The plots show a comparison of their expression levels in human (teal) and mouse (purple) across progenitor and signaling clusters.

**Fig. S7.**
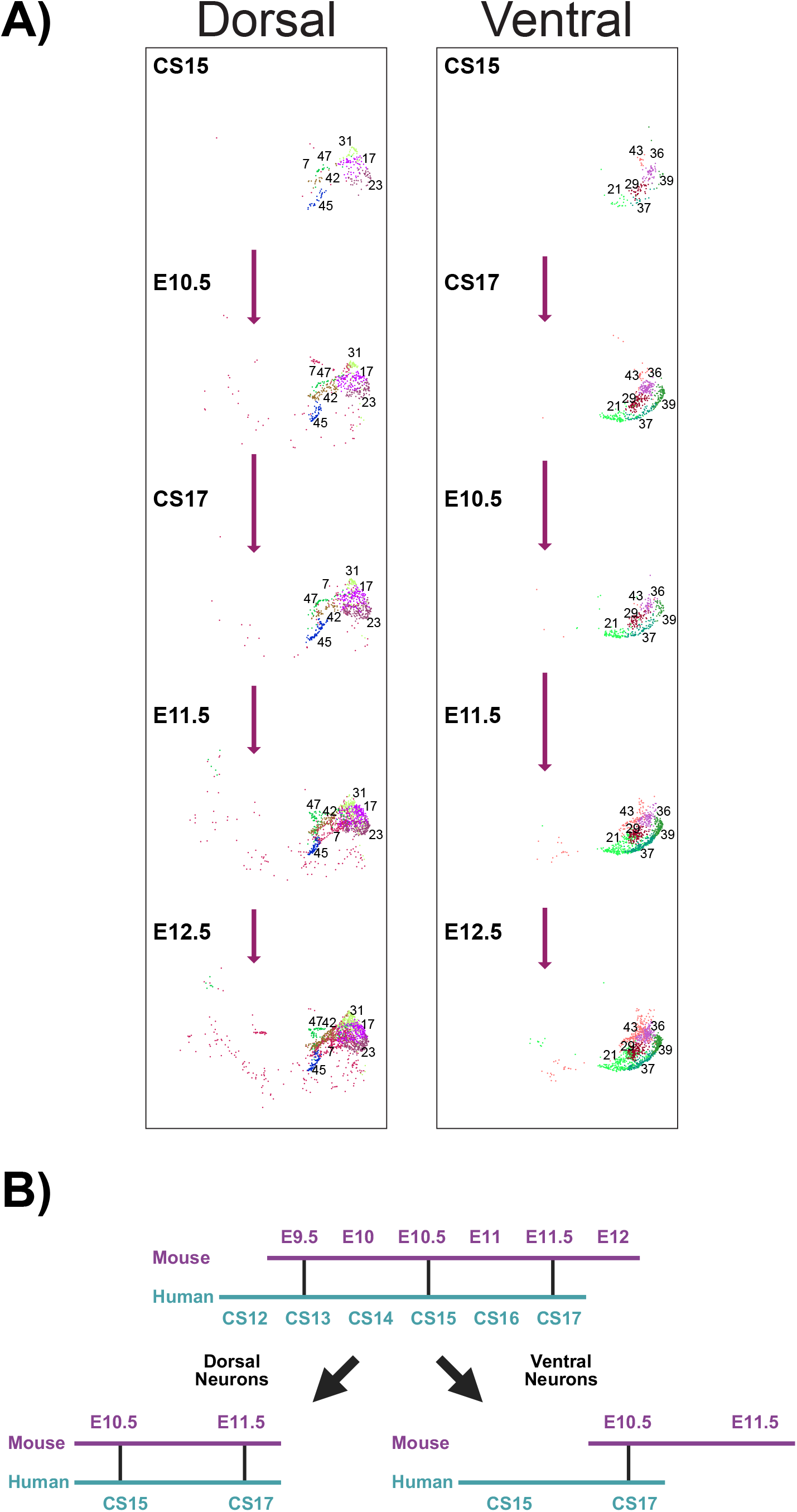
Cosine distances between human and mouse neuron populations separated by stage. **(A)** UMAP plots of integrated human and mouse dorsal (left) and ventral (right) neurons separated by stage and placed along a timeline from earliest to the latest stage. The length of arrows are proportional to the cosine distances between the stages. The clusters for each neural population are shown. **(B)** The equivalency between human and mouse ventral and dorsal neurons, showing a delay in ventral neurons, compared to the embryonic time line alignment.

## Bibliography

1. Li Wang, Cheng Wang, Juan A. Moriano, Songcang Chen, Guolong Zuo, Arantxa Cebrián-Silla, Shaobo Zhang, Tanzila Mukhtar, Shaohui Wang, Mengyi Song, Lilian Gomes de Oliveira, Qiuli Bi, Jonathan J. Augustin, Xinxin Ge, Mercedes F. Paredes, Eric J. Huang, Arturo Alvarez-Buylla, Xin Duan, Jingjing Li, and Arnold R. Kriegstein. Molecular and cellular dynamics of the developing human neocortex. Nature, pages 1–10, 2025. ISSN 0028-0836. doi: 10.1038/s41586-024-08351-7.

2. Vicki Metzis, Sebastian Steinhauser, Edvinas Pakanavicius, Mina Gouti, Despina Stamataki, Kenzo Ivanovitch, Thomas Watson, Teresa Rayon, S. Neda Mousavy Gharavy, Robin Lovell-Badge, Nicholas M. Luscombe, and James Briscoe. Nervous System Regionalization Entails Axial Allocation before Neural Differentiation. Cell, 175(4):1105–1118.e17, 2018. ISSN 0092-8674. doi: 10.1016/j.cell.2018.09.040.

3. Anna Kicheva and James Briscoe. Control of Tissue Development by Morphogens. Annual Review of Cell and Developmental Biology, 39(1), 2023. ISSN 1081-0706. doi: 10.1146/annurev-cellbio-020823-011522.

4. Rodrigo Senovilla-Ganzo and Fernando García-Moreno. The Phylotypic Brain of Vertebrates, from Neural Tube Closure to Brain Diversification. Brain, Behavior and Evolution, 99 (1):45–68, 2024. ISSN 0006-8977. doi: 10.1159/000537748.

5. Jan H. Lui, David V. Hansen, and Arnold R. Kriegstein. Development and Evolution of the Human Neocortex. Cell, 146(1):18–36, 2011. ISSN 0092-8674. doi: 10.1016/j.cell.2011.06.030.

6. Miki Nonaka-Kinoshita, Isabel Reillo, Benedetta Artegiani, Maria Ángeles Martínez-Martínez, Mark Nelson, Víctor Borrell, and Federico Calegari. Regulation of cerebral cortex size and folding by expansion of basal progenitors. The EMBO Journal, 32(13):1817–1828, 2013. ISSN 0261-4189. doi: 10.1038/emboj.2013.96.

7. Victor Borrell. Recent advances in understanding neocortical development. F1000Research, 8:F1000 Faculty Rev–1791, 2019. doi: 10.12688/f1000research.20332.1.

8. Colette Dehay and Wieland B. Huttner. Development and evolution of the primate neocortex from a progenitor cell perspective. Development, 151(4), 2024. ISSN 0950-1991. doi: 10.1242/dev.199797.

9. Alejandro L. Diaz and Joseph G. Gleeson. The Molecular and Genetic Mechanisms of Neocortex Development. Clinics in Perinatology, 36(3):503–512, 2009. ISSN 0095-5108. doi: 10.1016/j.clp.2009.06.008.

10. Jean M Hébert and Susan K McConnell. Targeting of cre to the Foxg1 (BF-1) Locus Mediates loxP Recombination in the Telencephalon and Other Developing Head Structures. Developmental Biology, 222(2):296–306, 2000. ISSN 0012-1606. doi: 10.1006/dbio.2000.9732.

11. Mattias Backman, Ondrej Machon, Line Mygland, Christiaan van den Bout, Weimin Zhong, Makoto M Taketo, and Stefan Krauss. Effects of canonical Wnt signaling on dorso-ventral specification of the mouse telencephalon. Developmental Biology, 279(1):155–168, 2005. ISSN 0012-1606. doi: 10.1016/j.ydbio.2004.12.010.

12. Ugo Borello and Alessandra Pierani. Patterning the cerebral cortex: traveling with morphogens. Current Opinion in Genetics & Development, 20(4):408–415, 2010. ISSN 0959-437X. doi: 10.1016/j.gde.2010.05.003.

13. H Bielen, S Pal, S Tole, and C Houart. Temporal variations in early developmental decisions: an engine of forebrain evolution. Current Opinion in Neurobiology, 42:152–159, 2017. ISSN 0959-4388. doi: 10.1016/j.conb.2016.12.008.

14. Arnaud Menuet, Alessandro Alunni, Jean-Stéphane Joly, William R. Jeffery, and Sylvie Rétaux. Expanded expression of Sonic Hedgehog in Astyanax cavefish:multiple consequences on forebrain development and evolution. Development, 134(5):845–855, 2007. ISSN 0950-1991. doi: 10.1242/dev.02780.

15. Arhat Abzhanov, Meredith Protas, B. Rosemary Grant, Peter R. Grant, and Clifford J. Tabin. Bmp4 and Morphological Variation of Beaks in Darwin’s Finches. Science, 305(5689):1462– 1465, 2004. ISSN 0036-8075. doi: 10.1126/science.1098095.

16. Ronald M. Bonett, Emily L. Bierbaum, Alexander J. Hess, Samantha D. Trame, Campbell W. Eckhardt, Madison A. Herrboldt, Carissa N. McGouran, Andrew D. Kolozsvary, Sydney Sowell, Ann Marie Flusche, and Rhiannon McGlone. Compounding heterochrony shapes the salamander visual system across adaptive zones. Proceedings B, 292(2055):20250910, 2025. doi: 10.1098/rspb.2025.0910.

17. Jonathan B. Sylvester, Constance A. Rich, Yong-Hwee E. Loh, Moira J. van Staaden, Gareth J. Fraser, and J. Todd Streelman. Brain diversity evolves via differences in patterning. Proceedings of the National Academy of Sciences, 107(21):9718–9723, 2010. ISSN 0027-8424. doi: 10.1073/pnas.1000395107.

18. J B. Sylvester, C A. Rich, C Yi, J N.Peres, C Houart, and J T. Streelman. Competing signals drive telencephalon diversity. Nature Communications, 4(1):1745, 2013. ISSN 2041-1723. doi: 10.1038/ncomms2753.

19. Kenneth J. McNamara. Heterochrony: the Evolution of Development. Evolution: Education and Outreach, 5(2):203–218, 2012. ISSN 1936-6426. doi: 10.1007/s12052-012-0420-3.

20. Karl Theiler. The house mouse: atlas of embryonic development. Springer-Verlag, New York, 1989. ISBN 0-387-05940-7.

21. Lorna Richardson, Shanmugasundaram Venkataraman, Peter Stevenson, Yiya Yang, Julie Moss, Liz Graham, Nicholas Burton, Bill Hill, Jianguo Rao, Richard A. Baldock, and Chris Armit. EMAGE mouse embryo spatial gene expression database: 2014 update. Nucleic Acids Research, 42(D1):D835–D844, 2014. ISSN 0305-1048. doi: 10.1093/nar/gkt1155.

22. Janet Kerwin, Yiya Yang, Paloma Merchan, Subrot Sarma, Jessica Thompson, Xunxian Wang, Juan Sandoval, Luis Puelles, Richard Baldock, and Susan Lindsay. The HUDSEN Atlas: a three-dimensional (3D) spatial framework for studying gene expression in the developing human brain. Journal of Anatomy, 217(4):289–299, 2010. ISSN 0021-8782. doi: 10.1111/j.1469-7580.2010.01290.x.

23. Dianne Gerrelli, Steven Lisgo, Andrew J. Copp, and Susan Lindsay. Enabling research with human embryonic and fetal tissue resources. Development, 142(18):3073–3076, 2015. ISSN 0950-1991. doi: 10.1242/dev.122820.

24. Christopher Y Chen, Nickesha C Anderson, Sandy Becker, Martin Schicht, Christopher Stoddard, Lars Bräuer, Friedrich Paulsen, and Laura Grabel. Examining the role of the surfactant family member SFTA3 in interneuron specification. PLOS ONE, 13(11):e0198703, 2018. doi: 10.1371/journal.pone.0198703.

25. Takeshi Shimizu and Masahiko Hibi. Formation and patterning of the forebrain and olfactory system by zinc-finger genes Fezf1 and Fezf2. Development, Growth & Differentiation, 51 (3):221–231, 2009. ISSN 1440-169X. doi: 10.1111/j.1440-169x.2009.01088.x.

26. Ugomma C. Eze, Aparna Bhaduri, Maximilian Haeussler, Tomasz J. Nowakowski, and Arnold R. Kriegstein. Single-cell atlas of early human brain development highlights heterogeneity of human neuroepithelial cells and early radial glia. Nature Neuroscience, 24(4): 584–594, 2021. ISSN 1097-6256. doi: 10.1038/s41593-020-00794-1.

27. Marc Fuccillo, Alexandra L. Joyner, and Gord Fishell. Morphogen to mitogen: the multiple roles of hedgehog signalling in vertebrate neural development. Nature Reviews Neuroscience, 7(10):772–783, 2006. ISSN 1471-003X. doi: 10.1038/nrn1990.

28. Elizabeth Manning, Kavitha Chinnaiya, Caitlyn Furley, Dong Won Kim, Seth Blackshaw, Marysia Placzek, and Elsie Place. Resolving forebrain developmental organisation by analysis of differential growth patterns. Nature Communications, 17(1):901, 2025. doi: 10.1038/s41467-025-67623-6.

29. Morgane Belle, David Godefroy, Gérard Couly, Samuel A. Malone, Francis Collier, Paolo Giacobini, and Alain Chédotal. Tridimensional Visualization and Analysis of Early Human Development. Cell, 169(1):161–173.e12, 2017. ISSN 0092-8674. doi: 10.1016/j.cell.2017.03.008.

30. Catherine Danesin, João N. Peres, Marie Johansson, Victoria Snowden, Amy Cording, Nancy Papalopulu, and Corinne Houart. Integration of Telencephalic Wnt and Hedgehog Signaling Center Activities by Foxg1. Developmental Cell, 16(4):576–587, 2009. ISSN 1534-5807. doi: 10.1016/j.devcel.2009.03.007.

31. Thomas Theil, Songül Aydin, Silke Koch, Lars Grotewold, and Ulrich Rüther. Wnt and Bmp signalling cooperatively regulate graded Emx 2 expression in the dorsal telencephalon. Development, 129(13):3045–3054, 2002. ISSN 0950-1991. doi: 10.1242/dev.129.13.3045.

32. Luca Muzio, Barbara Di Benedetto, Barbara DiBenedetto, Anastassia Stoykova, Edoardo Boncinelli, Peter Gruss, and Antonello Mallamaci. Emx2 and Pax6 Control Regionalization of the Pre-neuronogenic Cortical Primordium. Cerebral Cortex, 12(2):129–139, 2002. ISSN 1047-3211. doi: 10.1093/cercor/12.2.129.

33. Alexandra Gulacsi and Stewart A. Anderson. Shh Maintains Nkx2.1 in the MGE by a Gli3-Independent Mechanism. Cerebral Cortex, 16(suppl 1):i89–i95, 2006. ISSN 1047-3211. doi: 10.1093/cercor/bhk018.

34. Renée V. Hoch, John L.R. Rubenstein, and Sam Pleasure. Genes and signaling events that establish regional patterning of the mammalian forebrain. Seminars in Cell & Developmental Biology, 20(4):378–386, 2009. ISSN 1084-9521. doi: 10.1016/j.semcdb.2009.02.005.

35. Catherine Danesin and Corinne Houart. A Fox stops the Wnt: implications for forebrain development and diseases. Current Opinion in Genetics & Development, 22(4):323–330, 2012. ISSN 0959-437X. doi: 10.1016/j.gde.2012.05.001.

36. Jeremy A Cholfin and John L R Rubenstein. Patterning of frontal cortex subdivisions by Fgf17. Proceedings of the National Academy of Sciences, 104(18):7652–7657, 2007. ISSN 0027-8424. doi: 10.1073/pnas.0702225104.

37. Suoqin Jin, Maksim V. Plikus, and Qing Nie. CellChat for systematic analysis of cell–cell communication from single-cell transcriptomics. Nature Protocols, 20(1):180–219, 2025. ISSN 1754-2189. doi: 10.1038/s41596-024-01045-4.

38. Teresa Rayon, Despina Stamataki, Ruben Perez-Carrasco, Lorena Garcia-Perez, Christopher Barrington, Manuela Melchionda, Katherine Exelby, Jorge Lazaro, Victor L. J. Tybulewicz, Elizabeth M. C. Fisher, and James Briscoe. Species-specific pace of development is associated with differences in protein stability. Science, 369(6510):eaba7667, 2020. ISSN 0036-8075. doi: 10.1126/science.aba7667.

39. William D. Jones, Sarah M. Guadiana, and Elizabeth A. Grove. A model of neocortical area patterning in the lissencephalic mouse may hold for larger gyrencephalic brains. Journal of Comparative Neurology, 527(9):1461–1477, 2019. ISSN 0021-9967. doi: 10.1002/cne.24643.

40. Athéna R. Ypsilanti, Kartik Pattabiraman, Rinaldo Catta-Preta, Olga Golonzhka, Susan Lindtner, Ke Tang, Ian R. Jones, Armen Abnousi, Ivan Juric, Ming Hu, Yin Shen, Diane E. Dickel, Axel Visel, Len A. Pennacchio, Michael Hawrylycz, Carol L. Thompson, Hongkui Zeng, Iros Barozzi, Alex S. Nord, and John L. Rubenstein. Transcriptional network orchestrating regional patterning of cortical progenitors. Proceedings of the National Academy of Sciences, 118(51):e2024795118, 2021. ISSN 0027-8424. doi: 10.1073/pnas.2024795118.

41. Yutaka Maruoka, Norihiko Ohbayashi, Masamitsu Hoshikawa, Nobuyuki Itoh, Brigid L.M. Hogan, and Yasuhide Furuta. Comparison of the expression of three highly related genes, Fgf8, Fgf17 and Fgf18, in the mouse embryo. Mechanisms of Development, 74(1-2):175–177, 1998. ISSN 0925-4773. doi: 10.1016/s0925-4773(98)00061-6.

42. Stephen W Wilson and Corinne Houart. Early Steps in the Development of the Forebrain. Developmental Cell, 6(2):167–181, 2004. ISSN 1534-5807. doi: 10.1016/s1534-5807(04)00027-9.

43. Michael G.E. Goldschagg and Dorit Hockman. FGF18. Differentiation, 139:100735, 2024. ISSN 0301-4681. doi: 10.1016/j.diff.2023.10.003.

44. Jingyue Xu, Han Liu, Yu Lan, Bruce J. Aronow, Vladimir V. Kalinichenko, and Rulang Jiang. A Shh-Foxf-Fgf18-Shh Molecular Circuit Regulating Palate Development. PLoS Genetics, 12(1):e1005769, 2016. ISSN 1553-7390. doi: 10.1371/journal.pgen.1005769.

45. Thomas Theil, Elena Dominguez-Frutos, and Thomas Schimmang. Differential requirements for Fgf3 and Fgf8 during mouse forebrain development. Developmental Dynamics, 237(11):3417–3423, 2008. ISSN 1058-8388. doi: 10.1002/dvdy.21765.

46. Jeremy A. Cholfin and John L.R. Rubenstein. Frontal cortex subdivision patterning is coordinately regulated by Fgf8, Fgf17, and Emx2. Journal of Comparative Neurology, 509(2): 144–155, 2008. ISSN 0021-9967. doi: 10.1002/cne.21709.

47. Jyoti Rao, Zhisong He, Sebastian Loskarn, Audrey Bender, Youngmin Jo, Johanna Lückel, Martina Curcio, Irineja Cubela, J. Gray Camp, Barbara Treutlein, and Matthias P. Lutolf. Unified Generation of Regionalized Neural Organoids from Single-Lumen Neuroepithelium. bioRxiv, page 2025.11.18.689013, 2025. doi: 10.1101/2025.11.18.689013.

48. T. Fischer, J. Guimera, W. Wurst, and N. Prakash. Distinct but redundant expression of the Frizzled Wnt receptor genes at signaling centers of the developing mouse brain. Neuroscience, 147(3):693–711, 2007. ISSN 0306-4522. doi: 10.1016/j.neuroscience.2007.04.060.

49. Magnus Sandberg, Pierre Flandin, Shanni Silberberg, Linda Su-Feher, James D. Price, Jia Sheng Hu, Carol Kim, Axel Visel, Alex S. Nord, and John L.R. Rubenstein. Transcriptional Networks Controlled by NKX2-1 in the Development of Forebrain GABAergic Neurons. Neuron, 91(6):1260–1275, 2016. ISSN 0896-6273. doi: 10.1016/j.neuron.2016.08.020.

50. Oscar Marín. Development of GABAergic Interneurons in the Human Cerebral Cortex. European Journal of Neuroscience, 61(9):e70136, 2025. ISSN 0953-816X. doi: 10.1111/ejn.70136.

51. Longzhong Jia, Xiaohan Li, Yiming Yan, Linhe Xu, Jianbin Guo, Weichao Wang, Weirong Zhang, Lianyan Li, Borui Shang, Yiwei Zhang, Yashan Dang, Yuyan Zeng, Zhiyan Liao, Ruijuan Liang, Li Gu, Chenyi He, Zhen Long, Hanqing Hou, Yuhan Zhou, Mingchao Yan, Wei Huang, Lan Zhu, and Da Mi. Subventricular zone radial glial cells maintain inhibitory neuron production in the human brain. Science, 391(6782):eadw1803, 2026. ISSN 0036-8075. doi: 10.1126/science.adw1803.

52. Julien Delile, Teresa Rayon, Manuela Melchionda, Amelia Edwards, James Briscoe, and Andreas Sagner. Single cell transcriptomics reveals spatial and temporal dynamics of gene expression in the developing mouse spinal cord. Development, 146(12):dev.173807, 2019. ISSN 0950-1991. doi: 10.1242/dev.173807.

53. Yuhan Hao, Stephanie Hao, Erica Andersen-Nissen, William M. Mauck, Shiwei Zheng, Andrew Butler, Maddie J. Lee, Aaron J. Wilk, Charlotte Darby, Michael Zager, Paul Hoffman, Marlon Stoeckius, Efthymia Papalexi, Eleni P. Mimitou, Jaison Jain, Avi Srivastava, Tim Stuart, Lamar M. Fleming, Bertrand Yeung, Angela J. Rogers, Juliana M. McElrath, Catherine A. Blish, Raphael Gottardo, Peter Smibert, and Rahul Satija. Integrated analysis of multimodal single-cell data. Cell, 184(13):3573–3587.e29, 2021. ISSN 0092-8674. doi: 10.1016/j.cell.2021.04.048.

54. Anya Suppermpool, Chintan Trivedi, Gareth T. Powell, and Marta Andrés. Protocol for multiplex whole-mount RNA fluorescence in situ hybridization combined with immunohistochemistry in the mosquito brain. STAR Protocols, 6(4):104109, 2025. ISSN 2666-1667. doi: 10.1016/j.xpro.2025.104109.

55. Anika Klingberg, Anja Hasenberg, Isis Ludwig-Portugall, Anna Medyukhina, Linda Männ, Alexandra Brenzel, Daniel R. Engel, Marc Thilo Figge, Christian Kurts, and Matthias Gunzer. Fully Automated Evaluation of Total Glomerular Number and Capillary Tuft Size in Nephritic Kidneys Using Lightsheet Microscopy. Journal of the American Society of Nephrology, 28 (2):452–459, 2017. ISSN 1046-6673. doi: 10.1681/asn.2016020232.

56. Stuart Berg, Dominik Kutra, Thorben Kroeger, Christoph N. Straehle, Bernhard X. Kausler, Carsten Haubold, Martin Schiegg, Janez Ales, Thorsten Beier, Markus Rudy, Kemal Eren, Jaime I Cervantes, Buote Xu, Fynn Beuttenmueller, Adrian Wolny, Chong Zhang, Ullrich Koethe, Fred A. Hamprecht, and Anna Kreshuk. ilastik: interactive machine learning for (bio)image analysis. Nature Methods, 16(12):1226–1232, 2019. ISSN 1548-7091. doi: 10.1038/s41592-019-0582-9.

